# FOXM1 enhances DNA repair in aged cells to maintain the peripheral heterochromatin barrier to senescence enhancers

**DOI:** 10.1101/2025.10.29.685369

**Authors:** Carlos Sousa-Soares, Maria M. da Silva, José P. Castro, Joana C. Macedo, Elsa Logarinho

## Abstract

DNA damage is a key driver of aging, contributing to epigenetic erosion, senescence, and chronic inflammation. However, genoprotective strategies to counteract aging remain intangible. Here we show that FOXM1 repression during aging accounts for a global transcriptional shutdown of DNA repair genes and the accrual of DNA damage. Restored FOXM1 activity in aged cells reduces DNA damage and epigenetic alterations driving senescence. Mechanistically, FOXM1 drives the transcription of DNA repair genes, which prevents the DNA damage-driven degradation of the G9a methyltransferase and subsequent loss of H3K9me2 at the nuclear periphery. Remarkably, we show that amendment of the H3K9me2 guidepost for peripheral heterochromatin by FOXM1 induction in aged cells inactivates enhancers of the AP-1-driven senescence and inflammation program. These findings establish FOXM1 as an age-reversal factor capable of restoring (epi)genetic integrity to inhibit the senescence enhancer landscape, offering a promising therapeutic avenue to address the fundamental causes of aging.

## INTRODUCTION

Aging - the functional decline of organisms over time - is the highest risk factor for many prevalent human diseases (Jaul & Barron, 2017). Over the years, several features of aging at the molecular, cellular and physiological levels have been extensively characterized, being collectively known as the “hallmarks of aging” (López-Otín et al., 2023). Nevertheless, the main causal mechanisms of aging, as well as how different features of aging are hierarchically related, are still a subject of debate (de Magalhães, 2025; Gladyshev et al., 2024). Despite this, emergent evidence has pointed to DNA damage as a central driver of the aging process (Schumacher et al., 2021). For instance, many aged tissues accumulate multiple types of DNA damage, such as oxidative lesions or DNA double-strand breaks (Schumacher et al., 2021; Vijg, 2021; White & Vijg, 2016). In addition, somatic mutations accumulate over time due to errors in DNA repair (Cagan et al., 2022; Zhang et al., 2021). Interestingly, long-lived species have been found to display uniquely efficient DNA repair mechanisms, particularly regarding the repair of the highly harmful and mutagenic lesions such as DNA double-strand breaks (Lu et al., 2022; Ma et al., 2016; Tian et al., 2019; Tyshkovskiy et al., 2023). Other compelling evidence is that most human diseases of accelerated aging (i.e., progeroid syndromes) are caused by mutations in genes functionally linked to DNA repair or otherwise to nuclear stability (Schumacher et al., 2021). Finally, chronic exposure to DNA-damaging agents usually leads to premature aging, as observed in tissues targeted by chemo- and radiotherapy or skin exposed to ultraviolet radiation (Fisher Gary et al., 1997; Yousefzadeh et al., 2021).

These observations have raised the question of how structural changes in DNA ultimately cause aging phenotypes. Multiple evidence suggests that DNA damage is a strong driver of epigenetic erosion, which then leads to many downstream effects (Koch, Li, et al., 2025; Schumacher et al., 2021; Soto-Palma et al., 2022). For instance, pervasive cycles of DNA damage and repair may promote epigenetic alterations by relocating chromatin remodelers to damaged sites. This slowly erodes the epigenome over time, thus affecting normal gene expression and cell identity (Lu et al., 2023; Oberdoerffer et al., 2008; Yang et al., 2023). Moreover, persistent DNA damage evokes a set of signaling pathways known as the DNA damage response (DDR) (Jackson & Bartek, 2009), which is associated with profound changes in chromatin structure, including large-scale reorganization of nuclear compartments (Arnould et al., 2023; Dabin et al., 2023). The DDR also induces the transition into a senescent state, characterized by extensive chromatin remodeling and transcriptional changes dictating cell cycle arrest and adoption of a pro-inflammatory phenotype (Yousefzadeh et al., 2021; Zhang et al., 2020; Zhao et al., 2023). The accumulation of senescent cells gradually contributes to tissue decline over time, thus providing a bridge between DNA damage and organismal aging (López-Otín et al., 2023).

Assuming that DNA damage is at the root of the aging process, it is tempting to speculate that enhancing the expression of DNA repair genes represents a unified strategy to delay aging. In fact, such an approach could outcompete mainstream interventions, e.g. partial reprogramming and senolytics, that target aging phenotypes downstream of DNA damage (Guarente et al., 2024). However, targeting DNA repair has proven to be a challenge over the years. This is largely due to the complexity of most DNA repair pathways, which require a well-coordinated set of reactions involving multiple protein complexes acting in sequence (Bujarrabal-Dueso et al., 2025; Yousefzadeh et al., 2021). Therefore, it is most likely that targeting DNA damage requires the identification of a master regulator of DNA repair, which can coordinate the activity of many DNA repair factors at once.

In this work, we show that repression of the transcription factor *FOXM1* during aging is a key driver of DNA damage accrual, as well as of an ensuing epigenetic erosion at peripheral chromatin that drives a senescence transcriptional program. In human fibroblasts with advancing age, we found DNA repair pathways to be globally downregulated, concomitantly to increased levels of DNA damage. Downregulated DNA repair genes are enriched in targets of *FOXM1,* which together with B-MYB-MuvB (MMB) forms the master regulator complex of cell cycle-dependent gene expression (Engeland, 2018). We show that the downregulation of *FOXM1* leads to increased DNA damage, whose signaling triggers the proteasomal degradation of the G9a methyltransferase and subsequent global decrease in H3K9 dimethylation (H3K9me2) (Takahashi et al., 2012). Moreover, we uncovered that the loss of H3K9me2, an evolutionarily conserved epigenetic mark at the nuclear periphery (Poleshko et al., 2019), is prevalent at cis-regulatory regions of genes involved in inflammation and senescence, which we found to be transcriptionally upregulated following *FOXM1* repression. Notably, we demonstrated that ectopic expression of *FOXM1* in aged cells results in reduced DNA damage, recovery of H3K9me2 at the nuclear periphery, and rescue of senescence-associated phenotypes. These results disclose *FOXM1* as a master regulator of DNA repair, whose induction provides a unified intervention targeting DNA damage, epigenetic erosion, and senescence. Importantly, this provides mechanistic insight to our previous findings showing that an extra copy of *FOXM1* extends the maximum lifespan by 25% in both progeria and naturally aged mice (Ribeiro et al., 2022).

## RESULTS

### Age-associated accumulation of DNA damage in human fibroblasts

To confirm previous observations that DNA damage accumulates in aged cells (Schumacher et al., 2021; Vijg, 2021), we cultured human dermal fibroblasts (HDFs) derived from neonatal to octogenarian donors (Supplementary table 1). Cells were used at early passages, maintaining cumulative population doublings well below the limit for replicative senescence (Macedo et al., 2018). To detect DNA damage, we performed immunofluorescence against γH2AX and 53BP1 – two widely used biomarkers of DSBs (Kinner et al., 2008; Panier & Boulton, 2014) – in fibroblasts grouped in four age intervals. We observed that the percentage of nuclei positive for γH2AX or 53BP1 foci gradually increased with advancing age groups **(Fig. 1a, b)** and correlated positively with donors’ age (Supplementary Fig. 1a, b). Moreover, we observed that aged cells show a progressive accumulation of p53 and its target p21, consistent with an elevated DNA damage response (DDR) **(Fig. 1c, d)** (Engeland, 2022; Purvis et al., 2012).

**Figure 1.**
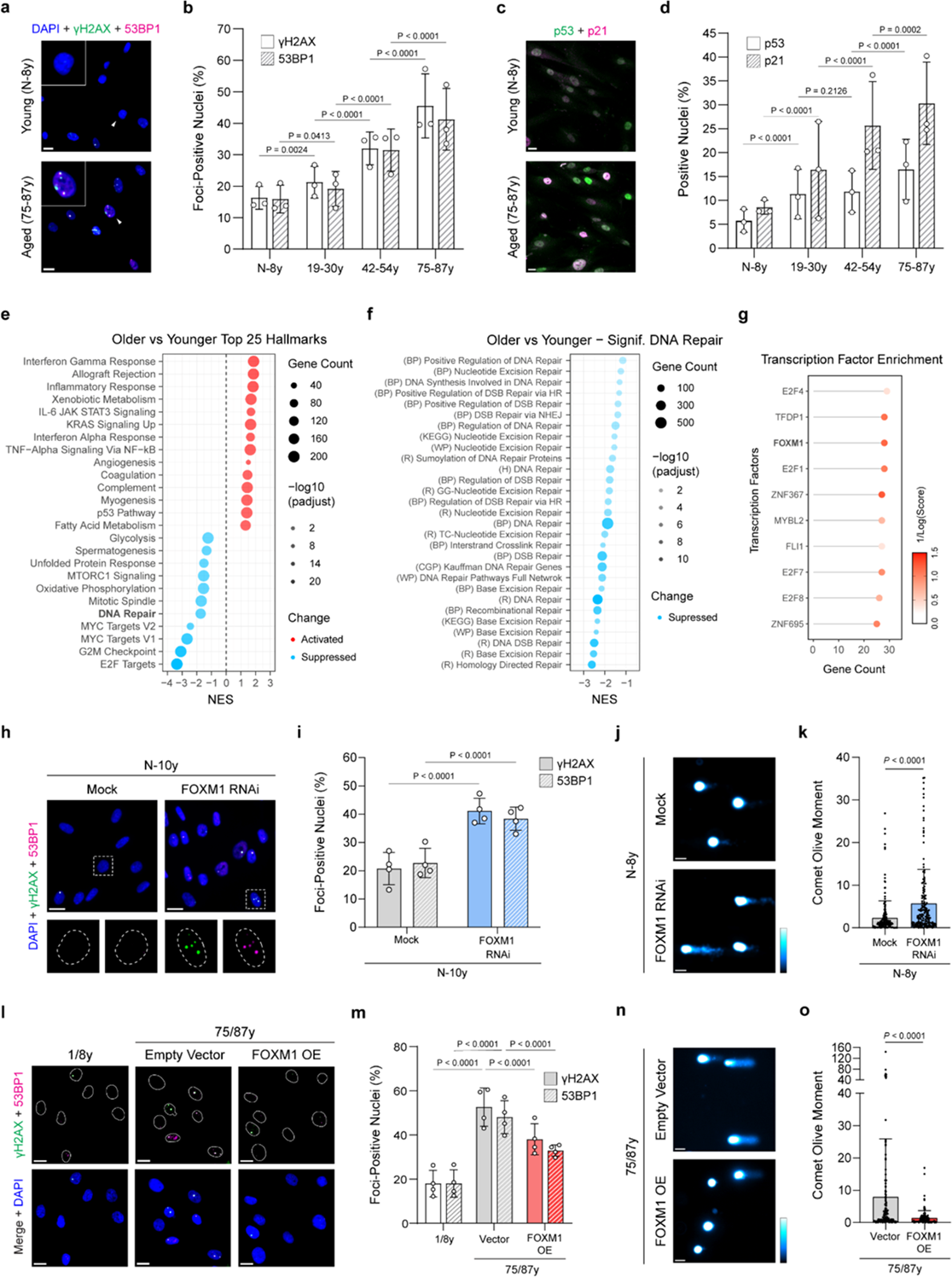
FOXM1 regulates DNA damage accrual in aging. **a**, Representative immunofluorescence images of γH2AX (green) and 53BP1 (magenta) in HDFs derived from young or aged donors. Arrowheads point to magnified nuclei in the insets. **b**, Percentage of nuclei positive for γH2AX or 53BP1 foci (DSBs markers) in HDFs from donors grouped by age intervals as indicated. Each dot represents a biological replicate of a distinct age (n = 400 cells per replicate). **c**, Representative immunofluorescence images of p53 (green) and p21 (magenta) in HDFs derived from young or aged donors. **d**, Percentage of cells staining positive for p53 or p21 (DDR markers) in HDFs from donors grouped by age intervals as indicated. Each dot represents a biological replicate of a distinct age (n = 400 cells per replicate). **e**, GSEA analysis comparing the older (>40-year-old) versus younger (1- to 40-year-old) cohorts using the “Hallmark” collection from MSigDB. Significant gene sets are ordered by normalized enrichment score (NES), and the top 25 terms ranked by *p*-adjust are displayed (see full list in Supplementary table 2). **f,** GSEA analysis comparing the older versus younger cohorts using gene sets related to DNA repair retrieved from the diverse collections available in MSigDB. Significantly enriched gene sets are ordered by normalized enrichment score (NES) (see also Supplementary table 3). **g**, Transcription factor enrichment analysis for downregulated DNA repair genes using ChEA3. Transcription factors are ordered by the number of gene associations. **h**, Representative immunofluorescence images of γH2AX (green) and 53BP1 (magenta) in young HDFs treated with either mock or FOXM1 RNAi. Arrowheads indicate nuclei shown in the insets, where dashed lines delimitate nuclear area. **i**, Percentage of nuclei positive for γH2AX or 53BP1 foci in mock or FOXM1 RNAi-treated HDFs. Each dot represents a biological replicate of a distinct age (n = 400 cells per replicate). **j**, Representative comet assay images and **k**, quantification of the comet olive moment in mock or FOXM1 RNAi-treated young HDFs (n ≥ 163 cells, 3 independent biological replicates of distinct age). Color scale used to interpret signal intensity is displayed on the right. **l**, Representative immunofluorescence images of γH2AX (green) and 53BP1 (magenta) in young or aged HDFs transduced with empty lentiviral vector or overexpressing *FOXM1* (FOXM1 OE). White lines delimitate nuclear areas. **n**, Percentage of nuclei positive for γH2AX or 53BP1 foci in HDF samples as mentioned in l). Each dot represents an independent experiment, using two biological replicates per group (n ≥ 350 cells per experiment). **m**, Representative comet assay images and **k**, quantification of the comet olive moment in aged HDFs transduced with empty lentiviral vector or overexpressing *FOXM1* (n ≥ 142 cells, 4 independent experiments using 2 biological replicates). Color scale used to interpret signal intensity is displayed on the right. Scale bars: 20 µm (**a**, **c**, **h**, and **i**) or 50 µm (**j** and **n**). Error bars indicate standard deviation. Statistics were performed comparing indicated groups using the Mann-Whitney test (**k**, **o**) and the two-tailed Fisher’s exact test (**b**, **d**, **i**, **m**).

Next, we evaluated whether the accumulation of DNA damage is caused by changes in gene expression related to DNA repair. To this end, we used public RNA-seq data of HDFs obtained from 133 donors of different ages (Fleischer et al., 2018) and performed differential gene expression analysis comparing an older cohort (> 40 years of age to nonagenarian) with a younger cohort (1- to 40-year-old). We found a clear separation of the two groups using principal component analysis (PCA) (Supplementary Fig. 1c) and 8339 genes to be differentially expressed (DEGs) (Supplementary Fig. 1d; Supplementary table 2). We then performed gene-set enrichment analysis (GSEA) (Subramanian et al., 2005) and interrogated the “hallmark” collection from MSigDB (Liberzon et al., 2015) **(Fig. 1e)**. The transcriptome of HDFs from aged donors displayed upregulation of signatures related to p53 activity and innate immune response (Tyshkovskiy et al., 2024; Tyshkovskiy et al., 2023; Varela et al., 2005), and downregulation of gene sets associated with proliferation and the cell cycle (Araújo et al., 2025; Macedo et al., 2018; Tomasetti et al., 2019). Interestingly, aged fibroblasts showed a downregulation of genes associated with DNA repair, in line with the observed accumulation of DNA damage. We further performed GSEA against several gene sets from MSigDB associated with DNA repair and specific DNA repair processes **(Fig. 1f)**. This analysis confirmed a global downregulation of DNA repair gene expression, as well as a significant repression of specific DNA repair processes. These include DSB repair pathways such as homologous recombination (HR) and non-homologous end joining (NHEJ) (Scully et al., 2019) (Supplementary Fig. 1e, f), but also other pathways involved in resolving bulky adducts (nucleotide excision repair - NER), simple base lesions such as oxidation (base excision repair – BER) or interstrand crosslinks (ICLs) (Supplementary Fig. 1g-i) (Huang & Zhou, 2021; Yousefzadeh et al., 2021). Finally, aggregation of HDFs into different age groups shows that differentially expressed DNA repair genes tend to become progressively downregulated during the lifespan (Supplementary Fig. 1j).

### FOXM1 repression contributes to DNA damage accrual in aging

To understand the key molecular players involved in the repression of DNA repair in aging, we obtained a list of genes downregulated in aged fibroblasts that are part of “Biological Processes: DNA Repair” – the largest downregulated gene set (Supplementary table 3). Next, we used ChEA3 (Keenan et al., 2019) to identify candidate transcription factors that are upstream regulators of those genes (**Fig. 1g**; Supplementary table 4). This analysis revealed an enrichment in several transcription factors that are well-known regulators of the DDR and DNA repair, such as E2F1 (Lin et al., 2001), the DREAM complex member E2F4 (Bujarrabal-Dueso et al., 2023; Chen et al., 2021), and FOXM1 (Zona et al., 2014). Interestingly, we found *FOXM1* itself to be downregulated in older cohort transcriptome (Supplementary Table 2). This is consistent with previous observations showing that *FOXM1* is downregulated in aged HDFs (Macedo et al., 2018) and murine tissues (Huang et al., 2023; Ribeiro et al., 2022). We thus hypothesized that *FOXM1* downregulation in aged cells may contribute to the accumulation of DNA damage.

To test this hypothesis, we evaluated the magnitude of DNA damage upon RNA interference (RNAi)-mediated knockdown of *FOXM1* in young HDFs (Supplementary Fig. 1k, l). We observed a significant increase in the percentage of nuclei positive for γH2AX and 53BP1 foci compared with the mock control, indicating an accumulation of unrepaired DSBs (**Fig. 1h, i**). We also performed the comet assay under alkaline conditions, which allowed the detection of both DSBs and other types of DNA lesions (e.g., interstrand crosslinks or single-strand lesions) (Olive & Banáth, 2006). This assay also revealed increased levels of DNA damage in young HDFs upon *FOXM1* knockdown (**Fig. 1j, k**). Moreover, *FOXM1* knockdown in young HDFs increased the percentage of nuclei positive for p53 or p21, consistent with elevated DNA damage levels (Supplementary Fig. 1m, n).

Next, we tested whether the DNA damage burden in elderly cells can be rescued by restoring FOXM1 levels. To this end, we ectopically expressed a constitutively active truncated form of *FOXM1* termed *FOXM1-dNdK* (Laoukili, Alvarez-Fernandez, et al., 2008; Laoukili, Alvarez, et al., 2008) in aged fibroblasts, under a doxycycline-inducible lentiviral system as previously reported (Ferreira et al., 2025; Macedo et al., 2018) (Supplementary Fig. 1o, p). FOXM1 overexpression in aged HDFs significantly reduced the proportion of nuclei positive for either γH2AX or 53BP1 foci in comparison with an empty vector infection control (**Fig. 1l, m**). FOXM1 overexpression also significantly reduced the global DNA lesion burden as measured by the alkaline comet assay (**Fig. 1n, o**). Finally, FOXM1 overexpression in aged HDFs decreased the percentage of nuclei positive for p53 or p21, consistent with an attenuation of the DNA damage response (Supplementary Fig. 1q, r).

### FOXM1 is a master regulator of DNA repair gene transcription in aging

To understand the molecular mechanisms by which FOXM1 regulates DNA repair in aging, we depleted *FOXM1* in young fibroblasts and overexpressed *FOXM1* in aged cells as described above. We found 3619 DEGs upon FOXM1 knockdown and 2187 DEGs upon ectopic *FOXM1* expression (Supplementary Fig. 2a, b) (Supplementary table 5), with PCA analysis showing a clear separation between conditions (Supplementary Fig. 2c). GSEA analysis revealed that, upon FOXM1 knockdown in young HDFs, DNA repair gene expression becomes significantly downregulated (**Fig. 2a, b**), as well as pathways involved in repairing specific types of damage (e.g., HR, NHEJ, NER, BER) (Supplementary Fig. 2d-f). Moreover, the ectopic expression of FOXM1 in aged cells resulted in a significant upregulation of DNA repair genes globally (**Fig. 2a, c**), as well as genes involved in specific DNA repair processes (Supplementary Fig. 2g-i). PCA of DNA repair gene expression revealed that FOXM1-depleted young fibroblasts cluster closely with aged HDFs, while FOXM1-overexpressing aged fibroblasts cluster with young controls (Supplementary Fig. 2j). Moreover, changes in DNA repair gene expression upon FOXM1 knockdown are positively correlated to those occurring in aged versus young fibroblasts and negatively correlated with those that occur after FOXM1 overexpression (Supplementary Fig. 2k). Genes downregulated by *FOXM1* knockdown or upregulated by *FOXM1* overexpression are also enriched in gene sets related to cell cycle and mitosis, in agreement with the well-established role of FOXM1 in regulating these processes (Supplementary Fig. 2l) (Fischer et al., 2022).

**Figure 2.**
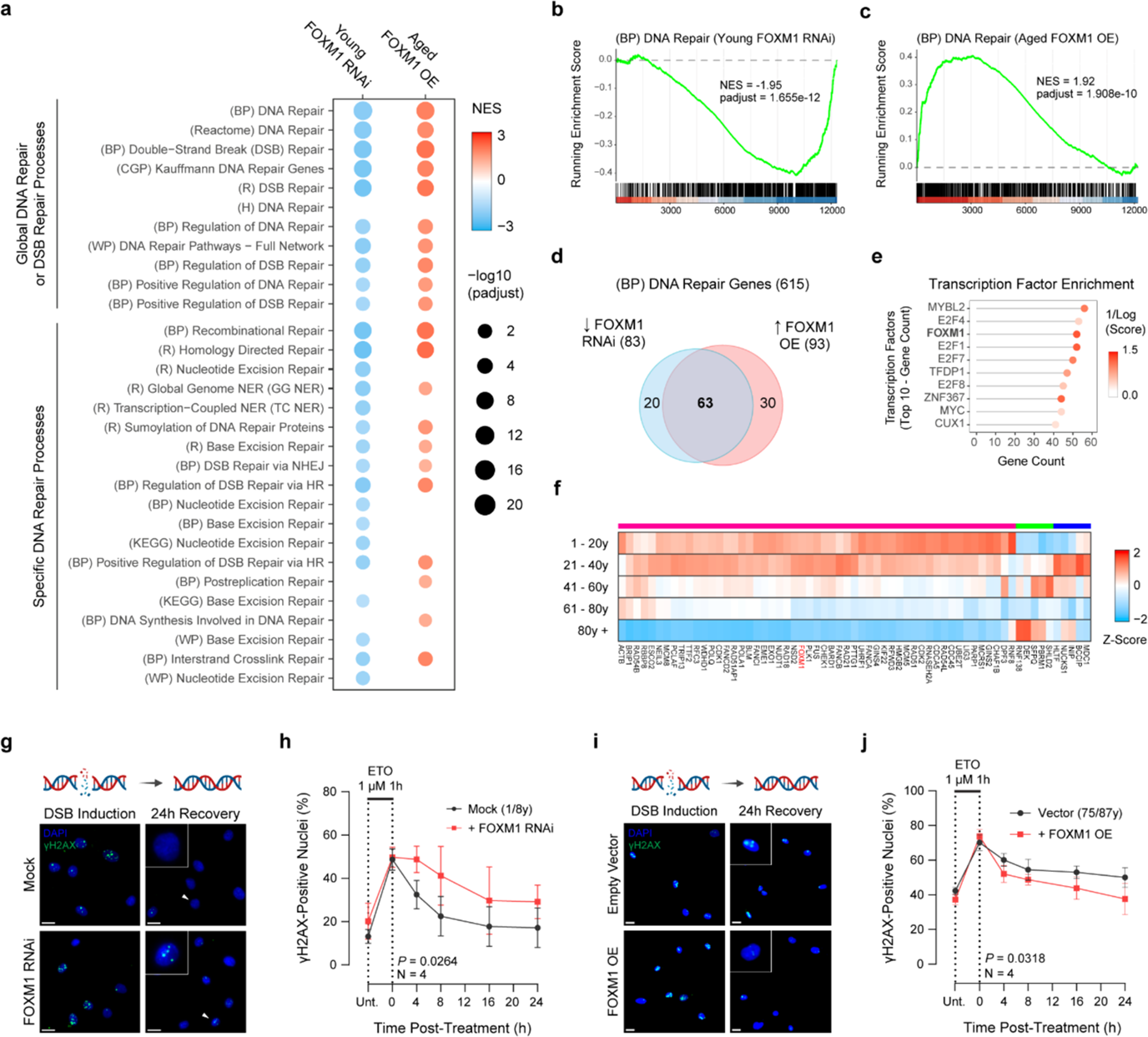
FOXM1 is a key regulator of DNA repair gene expression and increases DNA repair capacity. **a**, GSEA comparing FOXM1 RNAi versus mock treated young HDFs, and aged HDFs overexpressing *FOXM1* versus empty vector. Dots indicate significantly enriched gene sets related to DNA repair (MSigDB), divided into global and specific processes. For each condition, two biological samples were analyzed. **b, c**, GSEA enrichment plots showing downregulation of DNA repair (BP) genes upon FOXM1 knockdown in young HDFs (**b**), and upregulation of DNA repair (BP) genes upon FOXM1 OE in aged HDFs (**c**). **d**, Venn diagram displaying the intersection between robustly downregulated (Log2FC < -0.5, *p*-adjust < 0.05) DNA repair genes upon FOXM1 RNAi and robustly upregulated DNA repair genes (Log2FC > 0.5, *p*-adjust < 0.05) upon FOXM1 OE. The overlap is defined as the “FOXM1 DNA repair gene network”. **e**, Transcription factor enrichment analysis for the FOXM1 DNA repair gene network using ChEA3. Transcription factors were ordered by the number of gene associations. **f**, Heatmap showing a progressive z-scored mean expression (higher expression as red, lower expression as blue) of genes in the FOXM1 DNA repair network for human dermal fibroblasts grouped by different age intervals. *FOXM1* is highlighted in red. **g**, Representative images of γH2AX immunostaining (green) in young HDFs (mock or FOXM1 RNAi-treated) immediately after etoposide treatment and 24 hr following recovery. Arrowheads point to nuclei magnified in the insets. **h**, Percentage of nuclei staining positive for γH2AX in young HDFs, mock or *FOXM1* RNAi treated for 36 hr, that were then exposed to 1 µM etoposide for 1 hr and allowed to recover for the indicated hours (n = 4 independent experiments using HDFs from two distinct donors). **i**, Representative images of γH2AX immunostaining (green) in aged HDFs (empty vector or FOXM1 OE) immediately after etoposide treatment and 24 hr following recovery. Arrowheads point to nuclei magnified in the insets. **j**, Percentage of nuclei staining positive for γH2AX in aged HDFs transduced with empty vector or overexpressing *FOXM1* (FOXM1 OE) for 36h, that were then treated with 1 µM etoposide for 1 h and allowed to recover for the indicated hours (n = 4 independent experiments using HDFs from two distinct donors). Scale bars: 20 µm. Error bars represent the standard deviation. Statistics were performed comparing indicated groups using two-way analysis of variance (ANOVA) with Sidak’s multiple-comparison correction.

Next, we intersected DNA repair genes that were significantly downregulated upon FOXM1 knockdown and upregulated upon FOXM1 overexpression, resulting in a set of 63 genes robustly influenced by FOXM1 activity (**Fig. 2d**). Using ChEA3, we confirmed that these genes are enriched in candidate targets of FOXM1 and MYBL2 (B-MYB), both components of the MMB-FOXM1 activator complex (Fischer et al., 2022; Sadasivam et al., 2012) (**Fig. 2e**, Supplementary table 6). We further verified that most of the genes in this DNA repair network were progressively downregulated in HDFs retrieved from donors with advancing age (**Fig. 2f**). Network analysis shows that these genes cluster together mainly at the level of co-expression (Supplementary Fig. 3a), and are represented by well-known targets of FOXM1, such as *PARP1, RAD51, RAD21, CDK1*, *PLK1,* and others (Fischer et al., 2016; Sadasivam et al., 2012; Yu et al., 2023; Zona et al., 2014). Together, these results confirm the direct role of FOXM1 in driving DNA repair gene expression.

Interrogation of the *Tabula Muris Senis* public dataset (Almanzar et al., 2020) shows that, for more than 50 cell types, the expression of this network of DNA repair genes changes with aging *in vivo* (Supplementary Fig. 3b; Supplementary table 7). In agreement with transcriptomic data from HDFs, the vast majority of those cells showed a global downregulation of the network genes (e.g. trachea T-cells, bone marrow late pro-B cells – Supplementary Fig. 3c, d). This suggests that the FOXM1-dependent decline in DNA repair is a conserved feature of cellular aging *in vivo*.

To confirm that changes in DNA repair gene expression translate into a distinct DNA repair capacity, we knocked down FOXM1 in young fibroblasts and performed DNA-damage recovery assays using immunofluorescence. After short-term FOXM1 knockdown (36h), young HDFs were challenged with etoposide (1 µM, 1h) to induce DSBs and were allowed to recover for an additional 24h. FOXM1-depleted cells exhibited a significant delay in repairing DSBs, as indicated by a significantly higher proportion of γH2AX positive nuclei over time (**Fig. 2g, h**). Conversely, aged HDFs overexpressing *FOXM1* showed improved DSB repair kinetics versus control aged HDFs (**Fig. 2i, j**). We also tested if FOXM1 is required for NER of bulky lesions induced by UV-light, such as cyclobutane pyrimidine dimers (CPDs) (de Laat et al., 1999). Following UV-irradiation, the repair of CPDs over time was delayed in FOXM1-depleted cells, even at 24h post-irradiation (Supplementary Fig. 3e, f). In addition, we treated young HDFs with methyl methanesulfonate (MMS) – an alkylating agent whose damage is resolved mainly by BER (Sobol et al., 2002). FOXM1 RNAi-treated cells showed increased DNA damage levels after 24h, indicating decreased BER capacity (Supplementary Fig. 3g, h). Overall, these results confirm that changes in the DNA repair transcriptome driven by FOXM1 directly influence the ability of cells to repair their DNA.

### DNA damage-driven proteasomal degradation of the G9a methyltransferase leads to loss of H3K9me2-marked heterochromatin at the nuclear periphery during aging

Evidence suggests that DNA damage drives aging because it ultimately erodes the epigenetic landscape of cells (Lu et al., 2023; Siametis et al., 2021; Yang et al., 2023). Although epigenetic erosion appears particularly striking in heterochromatin-rich regions located at the nuclear periphery (López-Otín et al., 2023; Zhang et al., 2020), its mechanistic link with DNA damage is poorly understood. DNA damage signaling was shown to induce proteasomal degradation of G9a and GLP, major histone 3 lysine 9 mono- and dimethyltransferases, through p21-dependent activation of APC/C^Cdh1^ ubiquitin ligase. In turn, this causes a reduction in H3K9 dimethylation (H3K9me2) (Takahashi et al., 2012), a conserved mark of peripheral heterochromatin and lamina-associated domains involved in transcriptional silencing (Kind et al., 2013; Poleshko et al., 2019; Yokochi et al., 2009). Therefore, we hypothesized that DNA damage in aged cells may account for aging phenotypes by reducing G9a and H3K9me2 levels.

Using immunofluorescence, we confirmed that short-term (24h) induction of DNA DSBs by treating young HDFs with etoposide caused a pronounced reduction in both nuclear G9a and H3K9me2 signal intensities (**Fig. 3a, b**). This effect was significantly attenuated in the presence of the proteasome inhibitor MG132, confirming previous observations that DNA damage signaling triggers G9a proteasomal degradation (Takahashi et al., 2012). To test whether this mechanism is conserved across different types of genotoxic stress, we irradiated HDFs with UV light to induce bulky lesions. UV light led to a similar depletion of G9a and H3K9me2 after 24h (Supplementary Fig. 4a, b). In agreement with DNA damage accumulation along the lifespan, we found aged cells to display a significant reduction in G9a and H3K9me2 levels (**Fig. 3c, d**). These changes were correlated at the single-cell level (i.e., cells with lower levels of G9a tend to have lower levels of H3K9me2), thus reinforcing a functional relationship between G9a activity and heterochromatin maintenance (**Fig. 3e**). In addition, we used confocal microscopy to determine the preferential distribution of H3K9me2 at the nuclear periphery by quantifying i) the ratio of H3K9me2 fluorescence intensity between the nuclear periphery and interior (P/I ratio), and ii) the percentage of H3K9me2 fluorescence signal located at the nuclear periphery (see methods). Using this approach, we observed that the enrichment of H3K9me2 at the nuclear periphery becomes gradually compromised in HDFs as the donor age increases (**Fig. 3f-h**). These findings support a model in which age-associated DNA damage disrupts nuclear heterochromatin architecture by promoting the proteasome-mediated degradation of G9a, leading to the depletion of the repressive H3K9me2 mark at the nuclear periphery.

**Figure 3.**
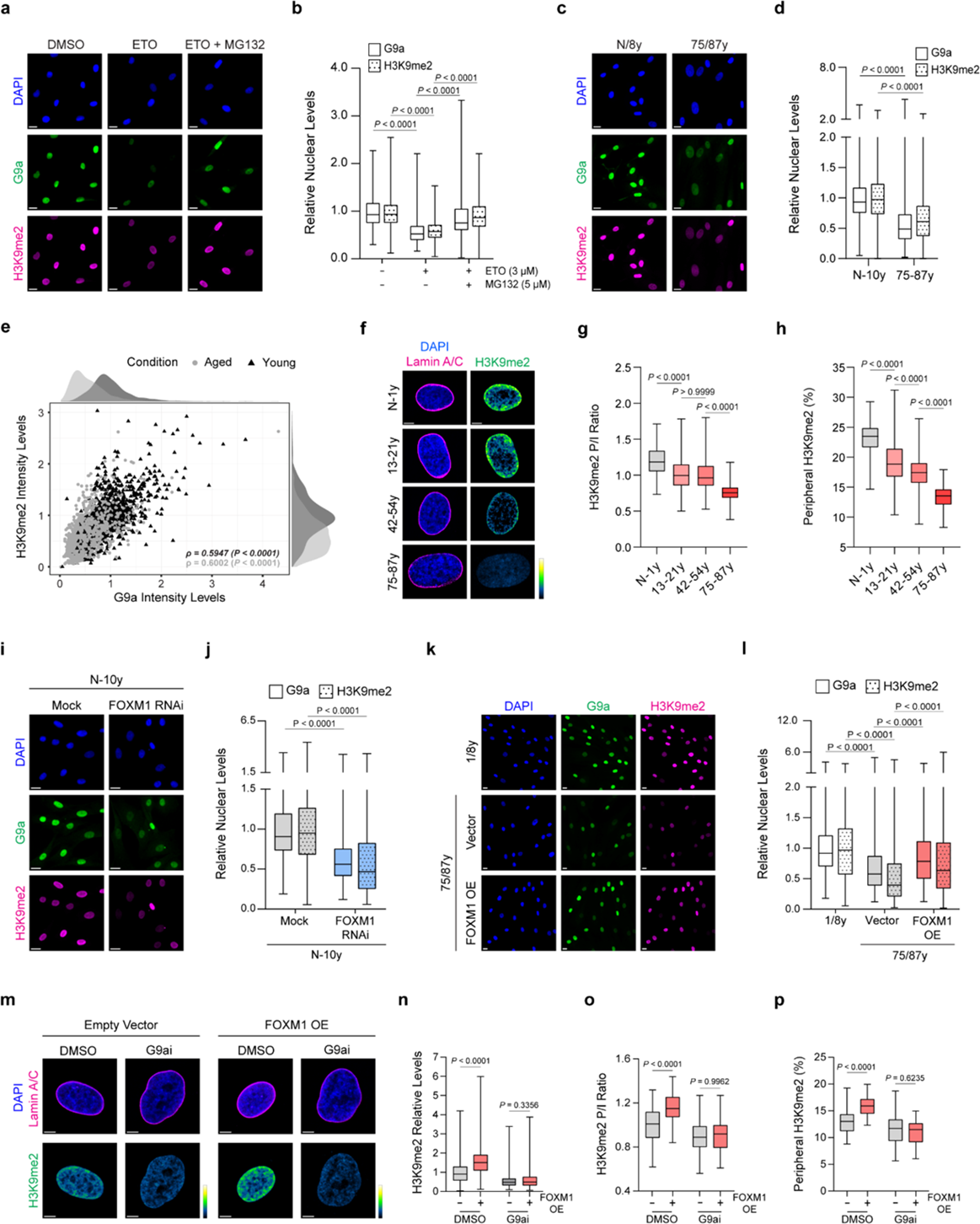
FOXM1 recovers G9a activity and H3K9me2 deposition at the nuclear periphery in aged cells by orchestrating DNA repair. **a,** Representative immunofluorescence images of G9a (green) and H3K9me2 (magenta) in young HDFs treated with either vehicle control (DMSO), etoposide (ETO 3 µM) or ETO in the presence of MG132 (5 µM; see methods for details). **b,** Quantification of G9a and H3K9me2 nuclear levels in young HDFs treated as mentioned in (a). Each condition is represented by n = 450 cells obtained from 3 independent biological replicates. **c,** Representative images of G9a (green) and H3K9me2 (magenta) immunostainings in young versus aged HDFs. **d,** Quantification of G9a and H3K9me2 nuclear levels in n = 900 cells obtained from 3 independent biological replicates per age condition. **e,** Single-cell correlation analysis between the fluorescence nuclear intensity levels of G9a and H3K9me2 in young and aged HDFs (see panels c, d). ρ = Spearman’s rank correlation coefficient. **f,** Representative confocal microscopy images of H3K9me2 (green-fire-blue gradient) and Lamin A/C (magenta) immunostainings in HDFs from donors grouped by age intervals as indicated (DNA in blue). **g, h,** Quantification of the H3K9me2 peripheral/internal signal brightness (P/I) ratio (**g**) and percentage of H3K9me2 signal at the nuclear periphery (**h**) in HDFs from donors grouped by age intervals as indicated (n ≥ 122 cells obtained from 3 independent biological replicates in each age group). **i,** Representative immunofluorescence images of G9a (green) and H3K9me2 (magenta) in young HDFs treated with mock vs. FOXM1 RNAi. **j,** Quantification of G9a and H3K9me2 nuclear levels in young HDFs treated with mock vs. FOXM1 RNAi. Each condition is represented by n = 1200 cells obtained from 4 independent biological replicates. **k,** Representative immunofluorescence images of G9a (green) and H3K9me2 (magenta) in young or aged HDFs transduced with empty lentiviral vector or overexpressing *FOXM1* (FOXM1 OE). **l,** Quantification of G9a and H3K9me2 nuclear levels in HDFs as mentioned in (k). Each group is represented by n = 1600 cells obtained from 4 independent experiments using 2 biological samples of different ages. **m,** Representative confocal microscopy images of H3K9me2 (green-fire-blue gradient) and Lamin A/C (magenta) immunostainings in aged HDFs overexpressing *FOXM1* vs. empty vector, in the presence of either DMSO or G9a inhibitor (BIX-01294). **n,** Quantification of H3K9me2 nuclear levels in aged HDFs as mentioned in (m). Each group is represented by n = 600 cells obtained from 3 independent replicates from 2 biological samples of different ages. **o, p,** Quantification of the H3K9me2 P/I ratio (o) and percentage of H3K9me2 signal at the nuclear periphery (**p**) in HDFs as mentioned in (m). Each group is represented by n ≥ 45 cells obtained from 3 independent biological replicates. Scale bars: 20 µm (**a, c, i** and **k**) or 5 µm (**f, m**). Statistics were performed comparing indicated groups using the Kruskal-Wallis test with Dunn’s multiple-comparison correction (**b, g, n, l**), the one-way ANOVA test with Sidak’s multiple-comparison correction (**h, o, p**), the Mann-Whitney test (**d, j**) and Spearman’s rank correlation (**e**).

### FOXM1 induction restores the H3K9me2 guidepost for peripheral heterochromatin in aged cells

Considering the striking changes in G9a and H3K9me2 with aging and their association with DNA damage, we hypothesized that FOXM1 activity may be important to preserve the H3K9me2 epigenetic guidepost during aging. In agreement with this, FOXM1 knockdown in young HDFs leads to a significant reduction in G9a and H3K9me2 levels (**Fig. 3I, j**), as well as in H3K9me2 enrichment at the nuclear periphery (Supplementary Fig. 4c-e). Conversely, the ectopic expression of FOXM1 in aged cells partially restored G9a and H3K9me2 levels (**Fig. 3k, l**), and the distribution of H3K9me2 near the nuclear lamina (Supplementary Fig. 4f-h). Next, we attempted to confirm that FOXM1 restores H3K9me2 specifically by restoring G9a protein levels. To this end, we overexpressed FOXM1 in aged HDFs in the presence of BIX 01294 - a specific G9ai inhibitor (G9ai) that disrupts H3K9me2 levels and its association with the nuclear lamina (Kind et al., 2013; Kubicek et al., 2007). The H3K9me2 levels in aged cells were further reduced in the presence of the G9ai, becoming almost completely depleted (**Fig. 3m**). Importantly, the G9ai blunted the effect of ectopic *FOXM1* expression in H3K9me2 resetting (**Fig. 3m-p**) indicating that FOXM1 induction restores H3K9me2 in a G9a-dependent manner.

Next, we performed CUT&RUN (Skene & Henikoff, 2017) against H3K9me2 using spike-in calibration to account for the observed differences in H3K9me2 global levels (Orlando et al., 2014). This allowed us to map differences in the genomic binding profile of H3K9me2 in young cells upon FOXM1 knockdown and aged cells upon FOXM1 overexpression. We observed that FOXM1 knockdown significantly decreased the binding of H3K9me2 to 14,103 genomic regions and FOXM1 overexpression increased H3K9me2 binding to 13,635 genomic regions (**Fig. 4a, b**; Supplementary Table 8). PCA analysis further demonstrated a clear separation between conditions (Supplementary Fig. 5a, b). In either case, we observed that differentially bound regions were enriched in previously annotated conserved lamina-associated domains (LADs) (Supplementary Fig. 5c) (Kind et al., 2015). This agrees with H3K9me2 being a hallmark of LADs (Kind et al., 2013; Poleshko et al., 2019) and with confocal microscopy data showing that H3K9me2 peripheral distribution is especially perturbed.

**Figure 4.**
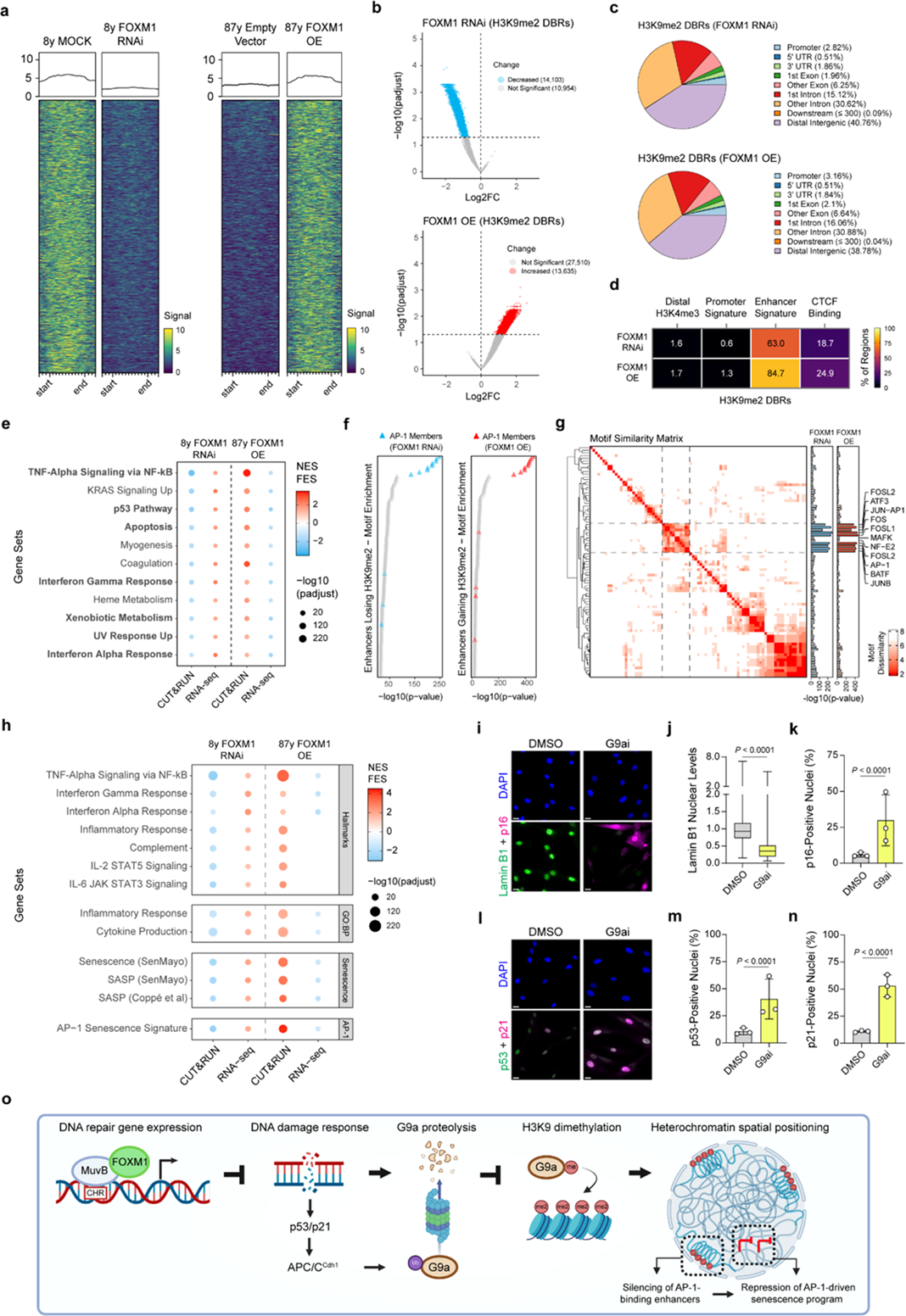
Maintenance of H3K9me2 levels by FOXM1 prevents the onset of senescence and inflammation-associated gene expression. **a**, Heatmap of the top thousand regions differentially bound by H3K9me2 (DBRs) upon *FOXM1* knockdown in young HDFs (mock vs. FOXM1 RNAi) or *FOXM1* overexpression in aged HDFs (empty vector vs. FOXM1 OE). Two biological replicates were used per condition. **b**, Volcano plots of H3K9me2 DBRs upon *FOXM1* knockdown in young HDFs and ectopic *FOXM1* expression in aged HDFs. **c**, Human genome annotation of H3K9me2 DBRs upon *FOXM1* knockdown in young HDFs and ectopic *FOXM1* expression in aged HDFs using ChIPSeeker (Yu et al., 2015). **d**, Heatmap showing the percentage of H3K9me2 DBRs in FOXM1 RNAi and FOXM1 OE overlapping with four main cCREs categories from ENCODE. **e**, Dot plot showing “Hallmark” gene sets inversely correlated between H3K9me2 GREAT analysis and RNA-seq GSEA analysis. Dots indicate gene sets enriched in regions losing H3K9me2 upon FOXM1 RNAi or gaining H3K9me2 upon FOXM1 OE in CUT&RUN data using GREAT, which are negatively correlated with RNA-seq data (i.e., upregulated upon FOXM1 RNAi and downregulated upon FOXM1 OE using GSEA). Color gradient indicates NES (for RNA-seq data) or the fold enrichment score FES (for CUT&RUN) (see Methods). Dot size indicates -log_10_ of adjusted *p*-value. **f**, Dot plot showing significantly enriched motifs (*q*-value < 0.05) for individual transcription factors in regions losing H3K9me2 upon FOXM1 RNAi or gaining H3K9me2 upon FOXM1 OE using HOMER. Triangles are ordered by nominal *p*-value and colored triangles indicate motifs targeted by known subunits of the AP-1 complex. **g**, Heatmap displaying the level of similarity between the top 100 motifs (*q*-value < 0.05, ranked by nominal *p*-value) enriched in enhancers losing or gaining H3K9me2 signal. Bar plots on the right indicate the enrichment *p-*value after log_10_ transformation. A cluster containing motifs targeted by several subunits of the AP-1 complex is highlighted using dashed lines. Transcription factors associated with each motif in HOMER are indicated on the right. **h**, Dot plot showing GSEA analysis in the CUT&RUN and RNA-seq datasets for “Hallmark” and “Biological Processes” related to inflammation and immunity, senescence and SASP gene sets curated and validated elsewhere, and the senescence gene signature associated with AP-1 activity (Martínez-Zamudio et al., 2020). Color gradient indicates NES (for RNA-seq data) or FES (for CUT&RUN). Dot size indicates -log_10_ of adjusted *p*-value. **i**, Representative images of Lamin B1 (green) and p16 (magenta) immunostainings in young HDFs treated either with G9a inhibitor (G9ai) or vehicle control (DMSO). **j**, **k**, Quantification of Lamin B1 nuclear levels (**j**) and p16-positive nuclei (**k**) in young HDFs treated as mentioned in (i). Each group is represented by n = 900 cells obtained from 3 independent biological replicates. **l**, Representative images of p53 (green) and p21 (magenta) immunostainings in young HDFs treated either with DMSO or G9ai. **m**, **n**, Quantification of p53-positive (**m**) and p21-positive (**n**) nuclei in young HDFs treated as mentioned in (l). Each group is represented by n = 900 cells obtained from 3 independent biological replicates. Scale bars: 20 µm (**i** and **l**). Statistics comparing indicated groups were performed using the Mann-Whitney test (**k**, **n**) and two-tailed Fisher’s exact (**b**, **j**, **m**), and standard methods for sequencing data (see Methods).

### FOXM1 maintains a peripheral chromatin landscape associated with cis-regulatory regions

To understand how modulation of peripheral heterochromatin by FOXM1 links DNA repair boosting to the recovery of youthful phenotypes, we characterized the genomic profile of H3K9me2 differentially bound regions (DBRs). Our analysis revealed that the differential binding of H3K9me2 upon FOXM1 knockdown or overexpression occurs mainly in non-coding regions, namely distal intergenic regions and introns (**Fig. 4c**). This agrees with the previously acknowledged repressive role of H3K9me2 in LADs and enhancers (Smith et al., 2021). Therefore, we hypothesized H3K9me2 DBRs to be over-represented in cis-regulatory regions, such as distal enhancers. Overlapping H3K9me2 DBRs with candidate cis-regulatory elements (cCREs) from ENCODE confirmed that they mostly exhibit an enhancer-like signature (63% in FOXM1 knockdown and 84.7% in FOXM1 overexpression), indicating that H3K9me2 levels regulate gene expression programs by modulating cCREs (**Fig. 4d**; Supplementary Table 9). Analysis of H3K9me2 DBRs using GREAT (Genomic Regions Enrichment of Annotations Tool) (McLean et al., 2010) showed that most regions-gene associations occur between 50-500 kb away from the transcription start site (TSS), in agreement with a cis-regulatory role (Supplementary Fig. 5d, e). GREAT was also used to perform functional enrichment of enhancers showing changes in H3K9me2. To consider the variation in H3K9me2 levels, we transformed the enrichment score (ES) of GREAT into a “fold enrichment score” (FES), where the enrichment of regions losing H3K9me2 was multiplied by -1 to yield a negative number, and the enrichment of regions gaining H3K9me2 was multiplied by 1 to yield a positive number (see Methods). Results show that enhancers losing H3K9me2 upon *FOXM1* knockdown or gaining H3K9me2 following *FOXM1* overexpression are associated with genes involved in mostly the same processes, but in opposite directions (Supplementary Fig. 5f). Many include inflammatory or stress responses (e.g. NF-κB activation, p53 pathway) and metabolic regulation (e.g. mTOR signaling).

### FOXM1 induction resets the H3K9me2-mediated inhibition of the AP-1-driven senescence program

The H3K9me2 histone mark is widely associated with transcriptionally repressed chromatin and its removal, unless compensated by other repressive mechanisms, may lead to transcriptional upregulation (Padeken et al., 2022). Thus, we focused on specific processes where i) loss of H3K9me2 upon FOXM1 RNAi correlated with transcriptional upregulation and ii) gain of H3K9me2 upon FOXM1 overexpression correlated with transcriptional silencing. To this end, we overlapped the GREAT enrichment with RNA-seq, maintaining gene sets in which H3K9me2 CUT&RUN enrichment (FES) and RNA-seq enrichment (NES) were opposite (**Fig. 4e**). This stringent approach revealed that enriched pathways were mostly related to innate immune signaling (e.g., “TNFα Signaling via NF-κB”, “Interferon Alpha Response”, “Interferon Gamma Response) and stress responses (e.g., “p53 Pathway”, “Apoptosis”).

In parallel, we performed motif enrichment analysis with HOMER (Heinz et al., 2010a) using as input the enhancers showing differences in H3K9me2 levels upon either *FOXM1* knockdown or overexpression. Sequence similarity-based clustering was used to obtain binding motifs that were consistently enriched. We found enhancers inside H3K9me2 DBRs to be enriched in binding motifs for members of the AP-1 heterodimer, such as FOS, JUNB and ATF3 (**Fig. 4f, g**; Supplementary table 10) (Bejjani et al., 2019). AP-1 is a master regulator of the cis-regulatory landscape of senescence, its associated secretory phenotype (SASP) and inflammatory signaling (Martínez-Zamudio et al., 2020), which are important hallmarks of aged cells (López-Otín et al., 2023). Motif enrichment analysis of the whole H3K9me2 DBRs also shows enrichment for motifs targeted by AP-1 subunits (Supplementary Fig. 5g; Supplementary table 11). This suggests that loss of H3K9me2 facilitates AP-1 binding to its target motifs (particularly at enhancers), and that H3K9me2 recovery by FOXM1 overexpression hinders this effect.

These results, combined with an enrichment for AP-1 binding motifs in H3K9me2-associated enhancers, led us to expand this analysis in the context of senescence and pro-inflammatory signaling. We therefore evaluated the enrichment of these enhancers in large gene sets related to inflammatory signaling from MSigDB (“Inflammatory Response” and “Cytokine Production”), well-characterized gene sets for senescence/SASP (e.g., SenMayo) (Coppé et al., 2008; Saul et al., 2022) and the AP-1 senescence gene signature (Martínez-Zamudio et al., 2020). All the senescence/inflammation gene sets were found to be upregulated upon FOXM1 RNAi and associated with enhancers losing H3K9me2, and conversely, to be downregulated upon ectopic FOXM1 expression and associated with enhancers gaining H3K9me2 (**Fig. 4h**). Focusing the analysis on all H3K9me2 DBRs (i.e., not exclusively associated with enhancers) led to similar results (Supplementary Fig. 5h). Importantly, the AP-1 gene signature of senescence (Martínez-Zamudio et al., 2020) was particularly enriched in enhancers showing changes in H3K9me2 occupancy, with most genes becoming upregulated upon *FOXM1* RNAi and downregulated upon FOXM1 ectopic expression (Supplementary Fig. 5i).

These observations suggest that H3K9me2 acts as gatekeeper of the senescence program. To test this, we treated young HDFs with the G9ai for 72h, to block deposition of H3K9me2. Treatment with the G9ai effectively reduced H3K9me2 levels and compromised its preferential distribution at the nuclear periphery (Supplementary Fig. 6a-e). Strikingly, this was sufficient to trigger several canonical senescence phenotypes (Ogrodnik et al., 2024), such as loss of Lamin B1 and an increase in p16-positive cells (**Fig. 4i-k**), as well as an increase in p53/p21 levels (**Fig. 4l-n**). Interestingly, other markers included the reduction of Ki-67-positive cells and accumulation of cells positive for γH2AX (Supplementary Fig. 6f-h). Overall, these results suggest that DNA-damage driven erosion of peripheral chromatin in aged cells unmasks an AP-1-driven senescence program, which can be offset by ectopic FOXM1 expression acting on DNA repair (**Fig. 4o**).

## DISCUSSION

DNA damage accumulates during the lifespan and is considered a key driver of aging (Schumacher et al., 2021; Vijg, 2021). Although increased DNA repair capacity has been linked to longevity, strategies capable of improving DNA repair have been missing due to the challenging complexity of the many pathways needed to repair distinct DNA damage insults (Bujarrabal-Dueso et al., 2025). Moreover, amongst a modest number of strategies reported to improve DNA repair capacity (Bujarrabal-Dueso et al., 2023; Francia et al., 2012; Mao et al., 2011; Oberdoerffer et al., 2008), only activation of NAD^+^-dependent enzymes such as sirtuins has been shown to robustly delay aging phenotypes in mammals (Bujarrabal-Dueso et al., 2025; Kanfi et al., 2012; Mitchell et al., 2014; Roichman et al., 2021; Satoh et al., 2013). Nevertheless, it is unclear whether such beneficial effects might be alternatively due to functions of sirtuins in epigenetic remodeling and/or metabolic reprogramming (Price et al., 2012; Ramadori et al., 2011; Roichman et al., 2021; Sundaresan et al., 2012; Van Meter et al., 2014). Therefore, it is of paramount interest to discover approaches that can promote healthy longevity by directly resetting global DNA repair capacity (Bujarrabal-Dueso et al., 2025).

Using cultured fibroblasts from different age donors, we confirmed that DNA damage gradually increases during the lifespan, consistent with previous observations across diverse tissues (Vijg, 2021; White & Vijg, 2016). Notably, we found fibroblasts from older age donors to display a consistent downregulation of genes involved in multiple DNA repair pathways, namely DSB repair, NER, BER and ICL repair, suggesting that DNA damage accrual correlates with a global repression of DNA repair genes. Transcription factors targeting repressed DNA repair genes include well-known master regulators of cell cycle gene expression, namely E2F1, E2F4, B-MYB and FOXM1 (Fischer et al., 2016). E2F1 drives early cell cycle gene expression, whereas FOXM1-MMB (a complex comprising FOXM1, B-MYB and the MuvB core) runs the expression of later cell cycle genes (Engeland, 2018). Contrariwise, E2F4, along with p107/p130 and the MuvB core, form the DREAM complex that outcompetes FOXM1-MMB and shuts down cell cycle gene expression (Fischer et al., 2016). Interestingly, DREAM activity was recently shown to inversely correlate with lifespan across species (Koch, Nandi, et al., 2025), and DREAM inhibition to enhance DNA repair in somatic cells (Bujarrabal-Dueso et al., 2023). Here, we show that increased FOXM1 activity restores DNA repair capacity in aged cells. The opposite activities of the DREAM and FOXM1-MMB complexes are thus the master regulators of global DNA repair capacity, whose targeting appears the logical approach to mitigate age-associated DNA damage.

We previously found that *FOXM1* expression is progressively shut down along aging (Macedo et al., 2018; Ribeiro et al., 2022) partly due to reduced activity of a distal enhancer that loses chromatin accessibility (Ferreira et al., 2025). Here, we disclose that *FOXM1* knockdown in young cells leads to overall repression of genes involved in key DNA repair pathways, ranging from DSB repair (e.g., HR, NHEJ) to NER and BER, thus mirroring the downregulation of DNA repair gene expression found in aged cells. Conversely, ectopic expression of *FOXM1* in aged cells increases DNA repair gene expression, thereby diminishing the accrual of DNA damage. Importantly, we found a DNA repair gene signature regulated by FOXM1 that includes several relevant players, such as *PARP1*, *CHK1, RAD21*, *RAD51*, *FANCA, FANCB* and *BLM* (Fischer et al., 2016; Yu et al., 2023; Zona et al., 2014). This signature is repressed in multiple cell types from diverse tissues during murine aging, indicating that *FOXM1* repression may explain DNA damage accrual in diverse tissues *in vivo*. Noteworthy, kinesin motors whose role in DSB DNA repair has been recently acknowledged, namely *KIF2C* and *KIF18B,* are FOXM1 transcriptional targets (Fischer et al., 2016; Luessing et al., 2021; Macedo et al., 2025; Zhu et al., 2020). Chemical enhancement of KIF2C activity was shown to reduce DSB DNA damage and to extend healthspan in the Hutchinson-Gilford progeria syndrome, highlighting the relevance of FOXM1 targets in genoprotection and aging (Macedo et al., 2025).

Besides DNA damage, aged cells display numerous epigenetic alterations, particularly loss of heterochromatin at the nuclear lamina (Liu et al., 2022; Zhang et al., 2020). Yet, how the accrual of DNA damage ensues these changes, and whether the improvement of DNA repair can reverse them, has remained elusive. One epigenetic alteration that occurs in direct response to DNA damage is the degradation of the G9a-GLP histone methyltransferase complex (Takahashi et al., 2012). G9a activity is essential to maintain H3K9me2 – an evolutionarily conserved histone mark of peripheral heterochromatin involved in gene repression (Poleshko et al., 2019; See et al., 2020; Tachibana et al., 2002). Our results show that *FOXM1* repression in aged cells causes a reduction of G9a and H3K9me2 levels (especially near the nuclear lamina), and that ectopic *FOXM1* expression restores G9a and H3K9me2 levels in aged cells. This suggests that *FOXM1*, by promoting DNA repair, restores the epigenetic landscape of peripheral chromatin that is lost with age. Interestingly, FOXM1 is also a negative regulator of FZR1/Cdh1 - a co-activator of the APC/C E3 ubiquitin ligase that is necessary for the proteolytic degradation of G9a (Takahashi et al., 2012) and whose levels we have found to increase with age (Araújo et al., 2025).

In addition, we assessed if the reversion of epigenetic changes in the nuclear lamina upon *FOXM1* induction was sufficient to rescue downstream hallmarks of aging, such as senescence and inflammation. We found that by restoring H3K9me2 levels at the nuclear periphery, FOXM1 prevents the onset of senescence and inflammation transcriptional programs. Mechanistically, we demonstrated that FOXM1 influences H3K9me2 occupancy predominantly at cis-regulatory regions, especially enhancers. This is in line with evidence showing that H3K9me2-modified enhancers become sequestered at the nuclear periphery to prevent their physical contact with unwarranted target genes (Fulmer et al., 2025; Smith et al., 2021). Enhancers inside regions losing H3K9me2 are predominantly enriched in binding motifs for several subunits of the AP-1 complex, a pioneer transcription factor of the senescence enhancer landscape (Martínez-Zamudio et al., 2020). Also, by integrating epigenomic with transcriptomic data, we found that loss of H3K9me2 leads to activation of enhancers associated with senescence and inflammation genes, which are also part of the AP-1 signature (Martínez-Zamudio et al., 2020). Interestingly, we also recently uncovered FOXM1 to negatively regulate chromatin accessibility at motifs targeted by AP-1 and to repress the promotor activity of several AP-1 subunits genes (Ferreira et al., 2025). Since AP-1 activity has also been shown to drive aging by hijacking organism maturation processes (Patrick et al., 2024), this opens the exciting possibility that FOXM1 may counteract developmental programs commandeered during aging.

Importantly, we envision the mechanistic link between DNA damage and senescence disclosed in this study to be conserved *in vivo*, given the observed repression of the FOXM1-DNA repair signature in multiple cell types during aging and the widespread nature of the H3K9me2 epigenetic mark. Along with the demonstration that long-term induction of a *FOXM1* transgene increases longevity in mice without tumorigenesis risk (Ribeiro et al., 2022), this study supports that FOXM1-mediated DNA repair and epigenetic maintenance plays a central role across many cell types during aging. By leveraging the genoprotective ability of FOXM1, we demonstrate a novel “bottom-up” approach for epigenetic rejuvenation, reminiscent of partial reprogramming but safer (Yücel & Gladyshev, 2024), senostatic rather than senolytic, therefore preserving beneficial senescence (Baker et al., 2023).

In conclusion, this work shows that ectopic expression of *FOXM1* is sufficient to restore DNA repair capacity and revert DNA damage-driven epigenetic erosion at the nuclear periphery that fosters senescence and loss of cell function/identity. These findings establish FOXM1 as an age reversal factor capable of restoring epigenetic integrity by enhancing DNA repair, offering a promising avenue to geroprotective therapies.

## METHODS

### Cell culture

Human dermal fibroblasts (HDFs) were derived from skin samples from apparently healthy neonatal to octogenarian donors (Zen Bio, NIGMS Human Genetic Cell Repository or Coriell Cell Repository). All details about the human fibroblasts used in this study are listed in Supplementary Table 1. Cells were cultured in Minimal Essential Medium (MEM) with Earle’s salts and L-glutamine (10-010-CV, Corning) supplemented with 15% Fetal Bovine Serum (FBS, from Gibco) and 1x Antibiotic-Antimycotic solution (Gibco). All cells were grown at 37 °C and in a humidified atmosphere with 5% CO_2_. Only early passage fibroblasts were used in all experiments (Barroso-Vilares et al., 2020; Macedo et al., 2018). For lentiviral production, HEK293T cells were cultured in Dulbecco’s modified Eagle medium (DMEM), supplemented with 10% FBS and 1x Antibiotic-Antimycotic (all from Gibco).

### RNA interference

HDFs were plated in serum-free culture medium and transfected with 50 nM of a small interfering RNA (siRNA) targeting FOXM1 (SASI_Hs01_00243977 from Merck). siRNA transfections were performed using Lipofectamine RNAiMAX (Thermo Fisher Scientific) in Opti-MEM medium (Gibco) according to the manufacturer’s instructions. Transfection medium was replaced by complete medium after 6 hr. Unless indicated, phenotypes were analyzed 72 hr following transfection.

### Lentiviral generation and transduction

Lentiviruses were produced using the Lenti-X Tet-ON Advanced Inducible Expression System (632162, Takara Bio). HEK293T helper cells cultured in serum-free medium were transfected with packaging plasmids pMd2.G (Addgene #12259) and psPAX2 (Addgene #12260) using Lipofectamine 2000 (Thermo Fisher Scientific) to generate lentiviruses carrying either pLVX–Tight-Puro (empty vector), pLVX–Tight-Puro–FOXM1-dNdK (Macedo et al., 2018), or pLVX–Tet-On Advanced, expressing the rtTA transactivator protein (Atze et al., 2016). Medium containing the lentiviruses was retrieved 24 hr after transfection, centrifuged and filtered to remove cell debris, and stored at -80 °C until further use. HDFs were infected for 8 hr with the transactivator lentiviruses, in combination with lentiviruses carrying either the empty vector or the FOXM1-dNdK construct. Transduction was carried out in a 1:1 ratio between transactivator and responsive lentivirus in the presence of 4 µg/mL polybrene and in serum-free medium. Induction of rtTA activity (and subsequent *FOXM1-dNdK* overexpression) was achieved by adding 500 ng/mL doxycycline to the medium. Unless indicated, phenotypes were analyzed 72 hr following doxycycline addition. Transfection efficiency was monitored by immunofluorescence.

### Immunofluorescence

Fibroblasts were grown on µ-Plate 24-well ibiTreat black plates (82406, ibidi) or on sterilized glass coverslips (2850-22, Corning) coated with 50 µg/mL fibronectin (F1141, Merck). Cells were fixed in freshly prepared 4% paraformaldehyde (PFA; 15713, EMS) in phosphate-buffered saline (PBS) for 10 min at room temperature (RT), washed 3 times with PBS, then permeabilized with 0.5% Triton X-100 for 10 min and washed 3 times with PBS-T (PBS with 0.05% Tween 20) for 5 min. Blocking was done with 10% FBS (Gibco) diluted in PBS-T for 1 hr at RT. Cells were incubated either 1h at RT or overnight at 4°C with primary antibodies diluted in PBS-T + 5% FBS as follows: rabbit anti-H3K9me2, 1:800 (39041, Active Motif); mouse anti-Lamin A/C, 1:100 (4777S, Cell Signaling); mouse anti-p21, 1:400 (clone F-5, sc-6246, Santa Cruz Biotechnology); rabbit anti-p53, 1:400 (clone FL-393, sc-6243, Santa Cruz Biotechnology); mouse anti-γH2AX, 1:1000 (clone JWB301, 05-636, Merck); rabbit anti-53BP1, 1:300 (4937, Cell Signaling Technology); mouse anti-G9a, 1:200 (clone A8620A, PP-A8620A-00, R&D Systems); mouse anti-Lamin B1, 1:400 (clone 3C10G12, 66095-1-Ig, Proteintech Group); rabbit anti-p16^INK4a^ 1:200 (ab7962, Abcam); rabbit anti-FOXM1, 1:500 (13147-1-AP, Proteintech Group); rabbit anti-Ki-67, 1:400 (ab15580, Abcam). Then, cells were washed 3 times with PBS-T for 5 min and incubated with secondary antibodies for 1 hr at RT - Alexa Fluor 488 (A11008, Thermo Fisher Scientific), Alexa Fluor 568 (A11031, Thermo Fisher Scientific) or Alexa Fluor 647 (A21244, Thermo Fisher Scientific) were diluted 1:1000 in PBS- T + FBS 5%. Samples were washed 2 times with PBS-T for 5 min and counterstained with DAPI (1 µg/mL; Merck) solution for 10 min at RT, then rinsed with PBS. Coverslips were mounted on slides (Epredia) using homemade mounting solution (90% glycerol, 0.5% N-propyl-gallate, 20 nM Tris, pH 8). For immunofluorescence of CPDs, the protocol was similar but with the following modifications: after permeabilization with 0.5% Triton X-100, cells were washed twice with PBS-T, once with PBS, and incubated with 2M HCl at RT for 30 min to denature DNA. Samples were then washed once with PBS, twice with PBS-T and blocked with 10% FBS to continue protocol as described above, using the following primary antibody: mouse hybridoma anti-CPDs, 1:500 (clone TDM-2, CAC-NM-DND-001, Cosmo-Bio).

The automated acquisition of images for quantification of protein global levels was performed using a Leica DMI6000 (Leica Microsystems, Germany) inverted epifluorescence microscope equipped with a Hamamatsu FLASH 4.0 camera (Hamamatsu, Japan) controlled by the LAS X v2.0 software. Images were acquired using either 20x/0.4NA or 40x/0.6NA dry objectives and appropriate filter cubes for each fluorophore. Confocal immunofluorescent images were taken under a Leica SP5 or SP8 laser scanning confocal microscope (Leica Microsystems, Germany) using a 63x/1.30NA glycerol objective controlled by the LAS AF (v2.6.0 software, Leica SP5) or LAS X (v3.5.6 software, Leica SP8) software. 2D images of interphase nuclei were taken as Z-stacks to reliably identify the equatorial plane of each nucleus. Identical settings were used for all matched images and minimum laser power was used to avoid signal saturation.

### Immunofluorescence image analysis

Image analyses were performed using ImageJ/Fiji software (Schindelin et al., 2012). For quantification of global protein levels, adjacent planes were stitched together with LAS X v3.7, and the DAPI signal was segmented using the “threshold” tool to automatically generate regions of interest (ROIs) corresponding to individual nuclei. Total fluorescence intensity was measured for each ROI as a measure of nuclear protein levels, after applying the background subtraction tool. For each independent experiment, data were normalized to the mean value of the control group. Cells positive for p16^INK4A^, p21 and p53 were automatically defined in an unbiased manner as having ≥ 2x fluorescence intensity of the mean of the corresponding control (young) population. Nuclei positive for γH2AX or 53BP1 were manually defined as having ≥ 1 bright, well distinguishable focus. Since cells in S-phase often accumulate pan-nuclear γH2AX or 53BP1 staining due to replication fork stalling and not necessarily DNA breaks (Ward & Chen, 2001), S-phase cells were detected using the DAPI fluorescence intensity (Roukos et al., 2015) and excluded from the analysis. For the quantification of CPD nuclear levels upon UV irradiation, CPD content was normalized to the DAPI signal (due to the stoichiometric relationship between DNA content and CPDs) (Bujarrabal-Dueso et al., 2023). This signal was then normalized to the maximum intensity in each condition, i.e., immediately following UV irradiation (0h).

To investigate the distribution of the H3K9me2 immunostaining signal at the nuclear periphery, we used the Lamin A/C signal to divide the nuclear equator into three compartments: the external perimeter of Lamin A/C (total nuclear area), the internal perimeter of Lamin A/C (nuclear interior area), and the area occupied by Lamin A/C itself (nuclear periphery area). To confirm the compartmentalization of H3K9me2, we used two distinct methods in which a higher value indicates higher compartmentalization: First, the mean gray value of H3K9me2 signal in the nuclear periphery was divided by the signal in the interior area (referred to as H3K9me2 P/I Ratio). Second, we divided the total H3K9me2 fluorescence intensity in the nuclear periphery by the total fluorescence intensity in the total nuclear area (% of peripheral H3K9me2) (Poleshko et al., 2019).

### Comet assay

Comet assays on HDFs were performed using a commercial kit (Comet Assay Kit, ab238544, Abcam) according to the manufacturer instructions, with small modifications. Briefly, cultured HDFs were detached from cell culture plates using TrypLE Express (Thermo Fisher Scientific), washed once with PBS, combined with comet agarose, transferred onto the top of an agarose base layer and maintained horizontally at 4 °C for 15 min. Slides were then transferred to pre-chilled lysis buffer in the dark for 1 hr and an alkaline solution for 30 min, at 4 °C. Alkaline electrophoresis was performed at 23 V for 30 min, after which slides were rinsed twice with distilled H_2_O and with 80% Ethanol for 5 min. Air dried slides were stained with the Vista Green DNA Dye at RT for 15 min. Images were performed under a Leica DMI6000 inverted epifluorescence microscope (Leica Microsystems) equipped with a Hamamatsu FLASH 4.0 camera (Hamamatsu, Japan) controlled by the LAS X v2.0 software, using a 20x/0.4NA objective. Images were analyzed with the Open Comet plugin v1.3.1 (Gyori et al., 2014) in ImageJ/Fiji, and the magnitude of DNA damage was quantified using the comet olive moment (percentage of DNA in the tail multiplied by the distance between the intensity-weighted centroids of head and tail) (Olive & Banáth, 2006).

### Drug treatments and UV irradiation

For the analysis of DNA repair capacity, HDFs were first subjected to either FOXM1 RNAi or FOXM1 OE for 36 hr. Then, cells were either a) washed twice with warm PBS 1x and irradiated with 40 J/m^2^ UV-C radiation (Vilber VL-6.LC) or b) incubated for 1 hr with complete medium containing 1 µM etoposide (ETO, S1225, 428 Selleck Chemicals) to induce DNA double-strand breaks (Burden & Osheroff, 1998). After washing twice with MEM, cells were incubated in complete medium and fixed in regular intervals up to an additional 24 hr using 4% PFA. To confirm proteasomal degradation of G9a upon DNA damage, young HDFs were either a) treated with 3 µM ETO for 28h or b) irradiated with 60 J/m^2^ UV-C, in the presence or absence of the proteasome inhibitor MG132 (5 µM) in the last 12h (see Takahashi et al., 2012). To examine the influence of G9a activity in H3K9me2 recovery, aged HDFs transduced with the *FOXM1-dNdK* transgene (versus empty vector) were treated with doxycycline for 72 hr to stimulate ectopic *FOXM1* expression, in the presence or absence of the specific G9a inhibitor BIX 01294 (G9ai, 3 µM) (Kubicek et al., 2007). To evaluate cellular phenotypes upon H3K9me2 reduction in young HDFs, cells were treated with BIX 01294 (G9ai, 1 µM) for 72 hr.

### RNA-seq analysis

Total RNA was extracted using Zymo’s Quick-RNA Microprep Kit (R1050, Zymo Research), according to manufacturer’s instructions. Paired-end raw sequencing reads were filtered by removing adapter sequences and nucleotides with low quality-score and aligned to human reference genome (GRCh38/hg38) using HISAT2 v2.21 (Kim et al., 2019). Aligned reads were processed with StringTie v2.2.3 (Pertea et al., 2015) to assemble reads into transcripts in the BMKCloud platform (www.biocloud.net). Public data using human dermal fibroblasts (Fleischer et al., 2018) were obtained by downloading counts matrices from NCBI-generated RNA-seq count data available from Gene Expression Omnibus (GEO). Differential expression analysis was performed using DESeq2 v1.42.1 (Love et al., 2014) from raw count matrices, with *p*-values for individual genes being corrected for multiple comparisons using the Benjamini-Hochberg (BH) method (Benjamini & Hochberg, 1995). PCA analysis was performed with DESeq2 for full transcriptome data after variance-stabilizing transformation, and with the factoextra package v1.0.7 (Lê et al., 2008) for DNA repair genes, to divide gene expression profiles into two k-means clusters, using transcripts per million (TPM) values as input. GSEA (Subramanian et al., 2005) was performed with the clusterProfiler package v4.16 in R (Wu et al., 2021), using gene lists ranked by Log2FC between conditions, gene sets and gene collections from MSigDB using the msigdbr package (v7.5.1) (Dolgalev, 2025) or custom gene sets indicated in the manuscript. Transcription factor enrichment analysis was performed using ChEA3 (Keenan et al., 2019). Since a lower score reflects higher enrichment, 1/log_10_ was applied to the score so that an increasing number reflects higher enrichment. The DNA repair correlation matrix was obtained using the corrplot package v0.95 (Hahsler et al., 2008) with Log2FC as input for each gene in the comparisons described. Network analysis of DEGs and gene-sets was performed using Metascape 3.5 (Zhou et al., 2019). Final networks were assembled using Cytoscape 3.10.2 (Shannon et al., 2003), filtering for the top ten significantly enriched clusters of biological processes. Heatmaps were obtained using ComplexHeatmap v2.24.1 (Gu et al., 2016) with z-scored TPM values for individual conditions or z-scored mean TPM values for each defined age group.

### *Tabula Muris Senis* data analysis

To investigate cell-specific gene expression of the FOXM1 DNA repair network during murine aging, we converted human genes to the corresponding mouse orthologs using g:Orth tool from g:Profiler (Kolberg et al., 2023). Next, we computed the network’s mean coefficient, using individual genes’ age-related coefficients from *Tabula Muris Senis* (Almanzar et al., 2020), and performed a one-sample t-test to assess whether the mean coefficient significantly deviated from zero in each cell type. Resulting *p*-values were adjusted for multiple testing using the Benjamini–Hochberg (BH) method, to control the false discovery rate, and then assigned a significance threshold of adjusted *p*-value (padjust) < 0.05. The top 35 cell types, ranked by significance and absolute effect size, were annotated directly on the plot using the ggplot2 package (v3.5.1) in R. To obtain representative UMAP plots, we analyzed pre-processed RDS objects from *Tabula Muris Senis* (https://cellxgene.cziscience.com/collections) with Seurat v5.2 (Hao et al., 2024) and attributed a global signature score to the FOXM1 network using the UCell package v2.12 (Andreatta & Carmona, 2021). Signature scores were then plotted alongside the age of the animal from which each cell was derived.

### CUT&RUN and next-generation sequencing

CUT&RUN was performed using the Active Motif ChIC/CUT&RUN Assay Kit (Active Motif, 53181) following manufacturers’ instructions, with small modifications. Briefly, 100.000 cells were detached from plates using TrypLE Express (Thermo Fisher Scientific) and then processed for nuclei isolation. Nuclei were attached to concanavalin A-bound magnetic beads, washed and incubated with primary antibodies overnight at 4 °C as follows: anti-H3K9me2 1:100 (39041, Active Motif), anti-rabbit IgG 1:100 (53181, Active Motif). After washing, nuclei were incubated with pAG-MNase fusion protein for 10 min at RT, and DNA cleavage induced for 2 hr by adding CaCl_2_ at 4 °C. *E. coli* spike-in DNA was added, and DNA fragments were isolated following incubation at 37 °C for 10 min. Illumina sequencing library was prepared using the Illumina TruSeq ChIP Library Preparation Kit according to the manufacturer’s instructions. The sequencing was performed with an Illumina NovaSeqX system.

### CUT&RUN data processing

Paired-end raw sequencing reads were pre-processed with fastp v1.0.1 (Chen et al., 2018) (quality profiling and adapter removal) and aligned to the human reference genome (GRCh38/hg38) using HISAT2 v2.2.1 (Kim et al., 2019) ignoring spliced alignment (--no-spliced-alignment). To avoid clonal artefacts, the duplicated mapped reads were removed using Samtools markdup v1.21 (Li et al., 2009). For spike-in control (Orlando et al., 2014) scale factor calculations, raw sequencing reads from exogenous spike-in DNA (Epicypher, 18-1401) were mapped to the *E. coli* reference genome (eschColi_K12), and scale factors between paired samples were calculated according to manufacturer’s instructions. Quality control and alignment were performed using the Galaxy platform (The Galaxy, 2024). The obtained BAM files were converted to bedGraph format using bedtools genomecov v2.31 (Quinlan & Hall, 2010) and peaks for each sample were called resorting to SEACR v1.3 (Sparse Enrichment Analysis for CUT&RUN) (Meers et al., 2019), using “stringent” mode and control (IgG) data to generate an empirical threshold. Differentially bound regions (DBRs) between conditions were obtained using DiffBind v3.16 (Ross-Innes et al., 2012; Stark & Brown, 2011), using aligned data for each sample and IgG control, together with peaks obtained with SEACR and scale factors. Changes in H3K9me2 occupancy between conditions (measured as Log2FC) were obtained using the “DBA_DESeq2” approach and obtained *p*-values corrected for multiple comparisons using the BH method. Regions were considered as differentially bound by H3K9me2 (DBRs) using a padjust threshold of < 0.05. Signal intensity heatmaps and PCA plots were generated with DiffBind. Overlap of H3K9me2 DBRs with genomic features was performed with ChIPSeeker v1.42.1 (Yu et al., 2015), using embedded features and previously annotated LADs from LAD Atlas (Kind et al., 2015). In the latter, *p*-values obtained using n = 5000 permutations. Overlap of H3K9me2 DBRs with regulatory elements from ENCODE was performed using bedtools intersect from the SCREEN registry (v3) (Abascal et al., 2020).

### GREAT analysis

DBRs in each condition (FOXM1 RNAi or OE) or the candidate enhancers overlapping with these elements, were used as input for rGREAT v2.8 (Gu & Hübschmann, 2023; McLean et al., 2010), resulting in enrichment score in specific gene sets. Gene sets from MSigDB (e.g., “Hallmarks”) were obtained using the msigdbr package (v7.5.1). We derived a new metric called “fold enrichment score” (FES), where the enrichment score was corrected by direction of changes in H3K9me2 occupancy: enrichment was multiplied by -1 for regions losing H3K9me2 (FES < 0), and enrichment was multiplied by 1 for regions gaining H3K9me2 (FES > 0).

### Motif enrichment analysis

Analysis of enriched motifs in H3K9me2 DBRs or candidate enhancers overlapping with these regions was performed using HOMER v4.11 (Heinz et al., 2010b), using the findMotifsGenome.pl tool. To obtain motif similarity matrices, consensus motifs from HOMER were used to build a matrix based on similarity between the sequence of characters for significant motifs, using the stringdist package v0.9.15 in R (Loo, 2014). Matrices were then plotted into a heatmap alongside log_10_-transformed *p*-values from HOMER using the ComplexHeatmap package.

### Statistical analysis

Sample sizes and statistical tests for each experiment are indicated in the figure legends or in the methods section. Normal distribution of the data was tested using the Shapiro–Wilk test. The appropriate statistical test (i.e., parametric versus non-parametric) was chosen according to data distribution. Unless specified, *p*-values were obtained using GraphPad Prism 8 (GraphPad). No statistical methods were used to predetermine sample sizes. Data collection and analysis were not performed blinded to the conditions of the experiments

## ACKNOWLEDGEMENTS

We thank Andrey Poleshko (UPenn, US) for providing valuable reagents and feedback on the manuscript. We thank the personnel at i3S Scientific Platforms for technical support: Genomics (A.M. Rocha, R. Mensink) and Advanced Light Microscopy (P. Sampaio, M. Azevedo, B. Monteiro). We thank D. Silva for providing resources for UV irradiation assays. We are grateful to all lab members for their helpful discussions.

## AUTHOR CONTRIBUTIONS

Conceptualization: EL, CSS. Data curation: CSS, JPC, MMS, JCM. Formal analysis: All authors. Investigation: All authors. Methodology: All authors. Visualization: All authors. Resources: EL. Funding acquisition: EL. Project administration: EL. Supervision: EL, JCM. Writing – original draft: CSS, EL. Writing – review & editing: CSS, EL.

## SUPPLEMENTARY INFORMATION

Supplementary Tables 1-11.

Supplementary Figures 1-6 and respective legends.

## DATA AVAILABILITY

Sequencing data will be deposited in the Gene Expression Omnibus database upon submission of the manuscript. All remaining materials are available from authors upon reasonable request.

## CORRESPONDENCE AND REQUEST FOR MATERIALS

Correspondence and requests for materials should be addressed to Elsa Logarinho.

