## Supplementary Figures for "FOXM1 enhances DNA repair in aged cells to maintain the peripheral heterochromatin barrier to senescence enhancers"

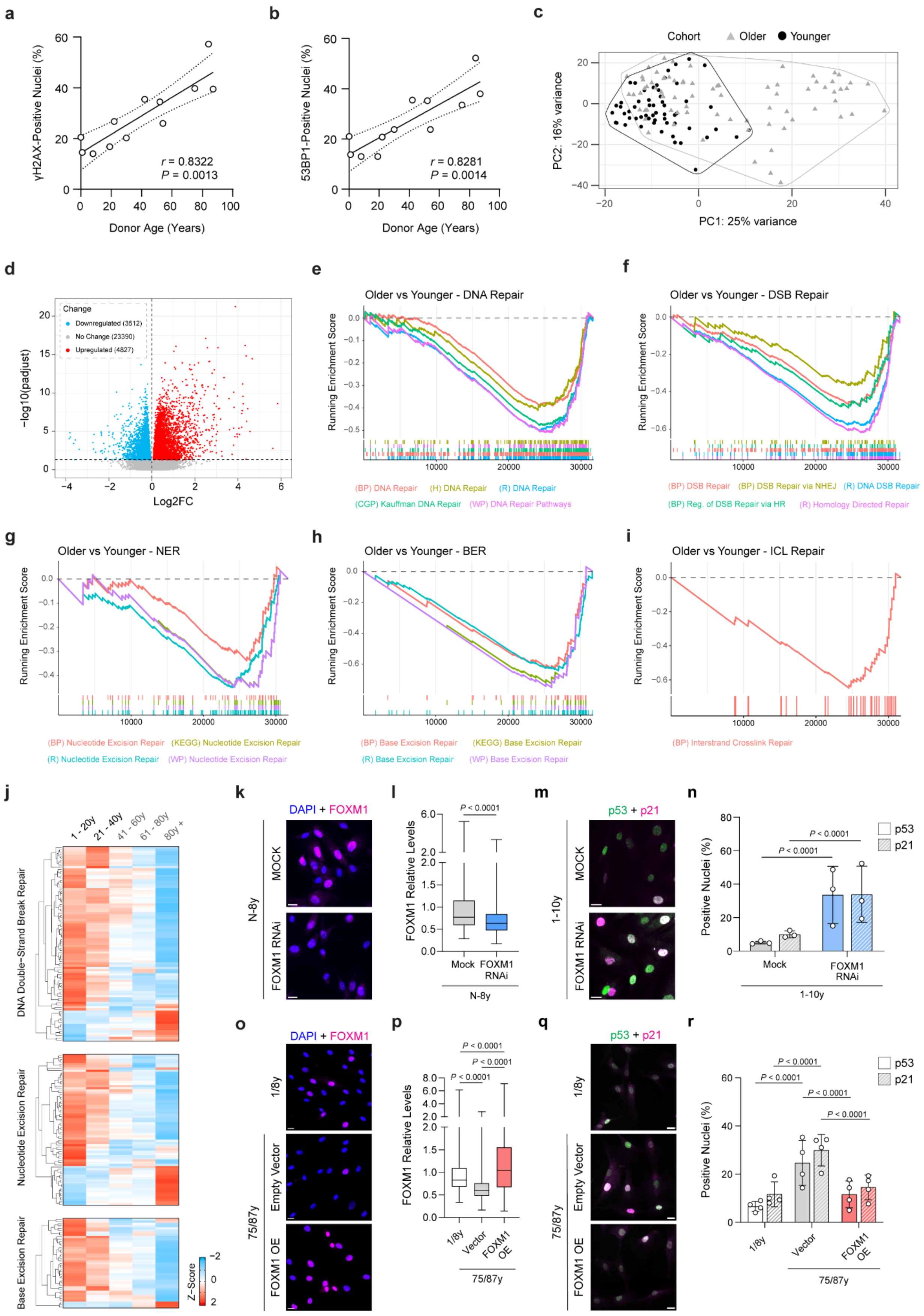


**SUPP. FIGURE 1**

**Supplementary Figure 1 – DNA repair gene repression in HDFs derived from donors with increasing age.** **a**, **b**, Correlation analysis between donor age and the percentage of nuclei positive for γH2AX (**a**) or 53BP1 (**b**) foci. ρ = Spearman's correlation coefficients. **c**, Principal component analysis (PCA) of the transcriptomes of HDFs retrieved from older (> 40-year-old) vs. younger (< 40-year-old) donors. **d**, Volcano plot of changes in gene expression between HDFs from younger vs. older cohorts (GSE113957). **e-i**, GSEA plots showing downregulation of gene sets related to DNA repair (**e**), DSB repair including HR and NHEJ (**f**), NER (**g**), BER (**h**) and ICL repair (**i**) in older vs. younger HDFs. **j**, Heatmap showing a progressive z-scored mean expression (higher expression as red, lower expression as blue) of DEGs belonging to the "DNA Double Strand Break Repair", "Nucleotide Excision Repair" and "Base Excision Repair" from the Reactome subcollection from MSigDB. **k**, Representative immunofluorescence images of FOXM1 (magenta) in young HDFs treated with mock or FOXM1 RNAi. **l**, Quantification of FOXM1 nuclear protein levels in HDFs treated as mentioned in (k). Each group is represented by n = 1500 cells from 3 independent biological replicates of different ages. **m**, Representative immunofluorescence images of p53 (green) and p21 (magenta) in young HDFs treated with mock or FOXM1 RNAi. **n**, Quantification of the percentage of nuclei positive for p53 and p21 in HDFs treated as mentioned in (m). Each dot represents a biological replicate of distinct age per condition (n = 300 cells per replicate). **o**, Representative immunofluorescence images of FOXM1 (magenta) in young or aged HDFs transduced with empty lentiviral vector or overexpressing *FOXM1* (FOXM1 OE). **p**, Quantification of FOXM1 nuclear protein levels in HDFs treated as mentioned in (o). Each group is represented by n = 1600 cells from 4 independent experiments using 2 biological replicates of different ages. **q**, Representative immunofluorescence images of p53 (green) and p21 (magenta) in young or aged HDFs transduced with empty lentiviral vector or overexpressing *FOXM1* (FOXM1 OE). **r**, Percentage of nuclei positive for p53 and p21 in HDFs treated as mentioned in (q). In each condition, each dot represents an independent experiment from a total of n = 4 using two distinct biological samples (n = 300 cells quantified per replicate). Scale bars: 20 μm. Error bars represent the standard deviation. Statistics comparing indicated groups were performed using the Mann-Whitney test (**l**), Kruskal-Wallis test with Dunn's multiple-comparison correction (**p**), and the two-tailed Fisher exact test (**n**, **r**). Correlation analyses were performed using Spearman's rank correlation test (**a**, **b**). Statistics for sequencing data are described in detail in Methods.

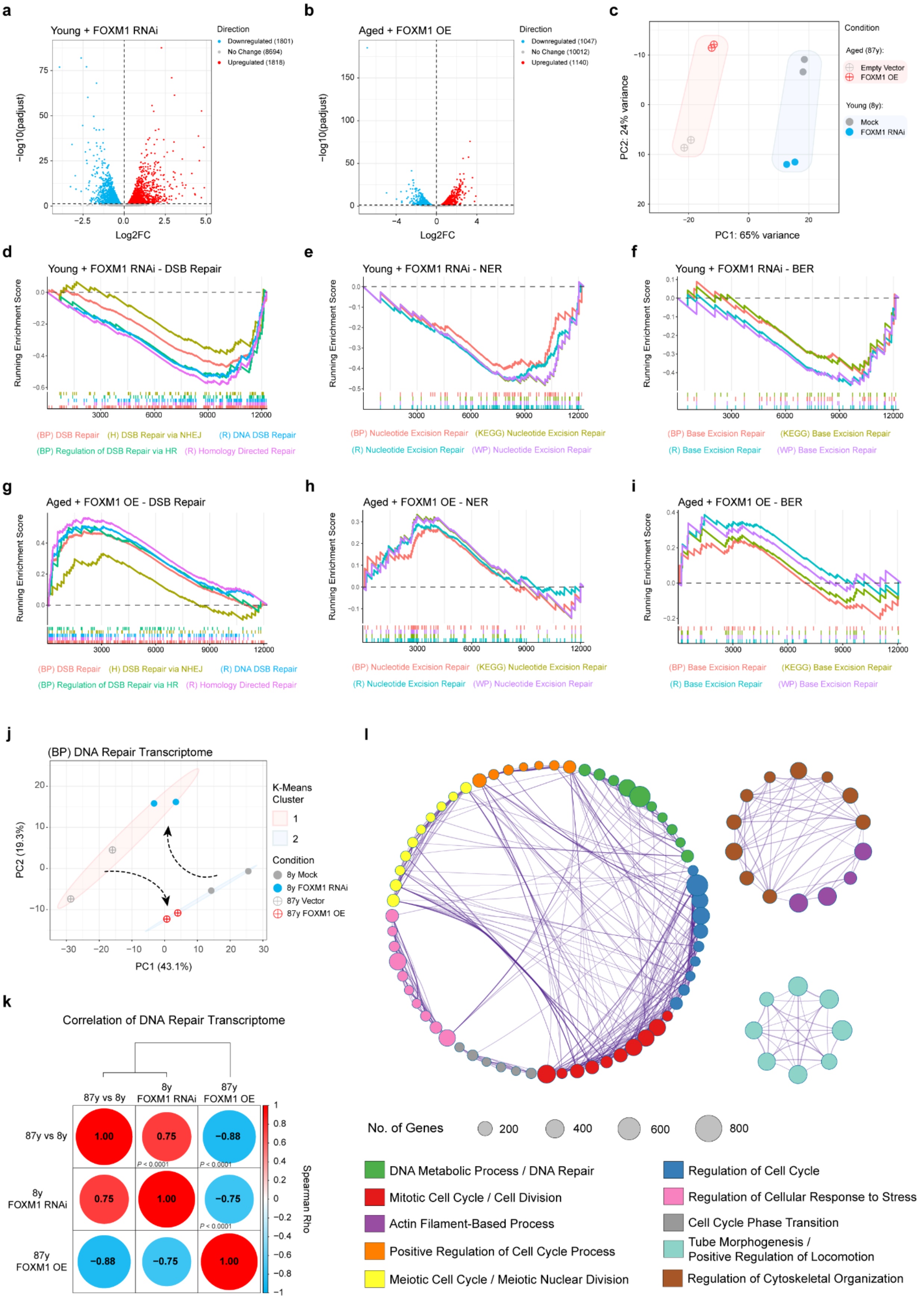


37

**SUPP. FIGURE 2**

**Supplementary Figure 2 – FOXM1 counteracts age-associated changes in the expression of DNA repair genes related to multiple pathways.** **a**, **b**, Volcano plots showing Log2FC versus -$\log_{10}$(padjust) values for individual gene expression upon *FOXM1* knockdown in young HDFs (**a**) or ectopic *FOXM1* expression in aged HDFs (**b**). **c**, PCA plot of young HDFs treated with either mock or FOXM1 RNAi, and aged HDFs transduced with empty vector or overexpressing *FOXM1*. **d-f**, GSEA plots showing downregulation of gene sets related to DSB repair (**d**), NER (**e**), BER (**f**) upon *FOXM1* knockdown. **g-i**, GSEA plots showing upregulation of gene sets related to DSB repair (**g**), NER (**h**), BER (**i**) upon ectopic *FOXM1* expression. **j**, PCA plot of DNA repair genes (Gene Ontology - BP) of young HDFs treated with either mock or FOXM1 RNAi, and aged HDFs transduced with empty vector or overexpressing FOXM1, coupled with k-means clustering. **k**, Correlogram showing the correlation between changes in DNA repair gene expression (Gene Ontology - BP) using Log2FC for the indicated comparisons. Spearman rank correlation coefficients (Spearman rho) are displayed inside the circles. **l**, Network of enriched terms for genes downregulated by *FOXM1* knockdown and upregulated by *FOXM1* overexpression using Metascape (Zhou et al., 2019). Statistics for sequencing data are described in detail in Methods.

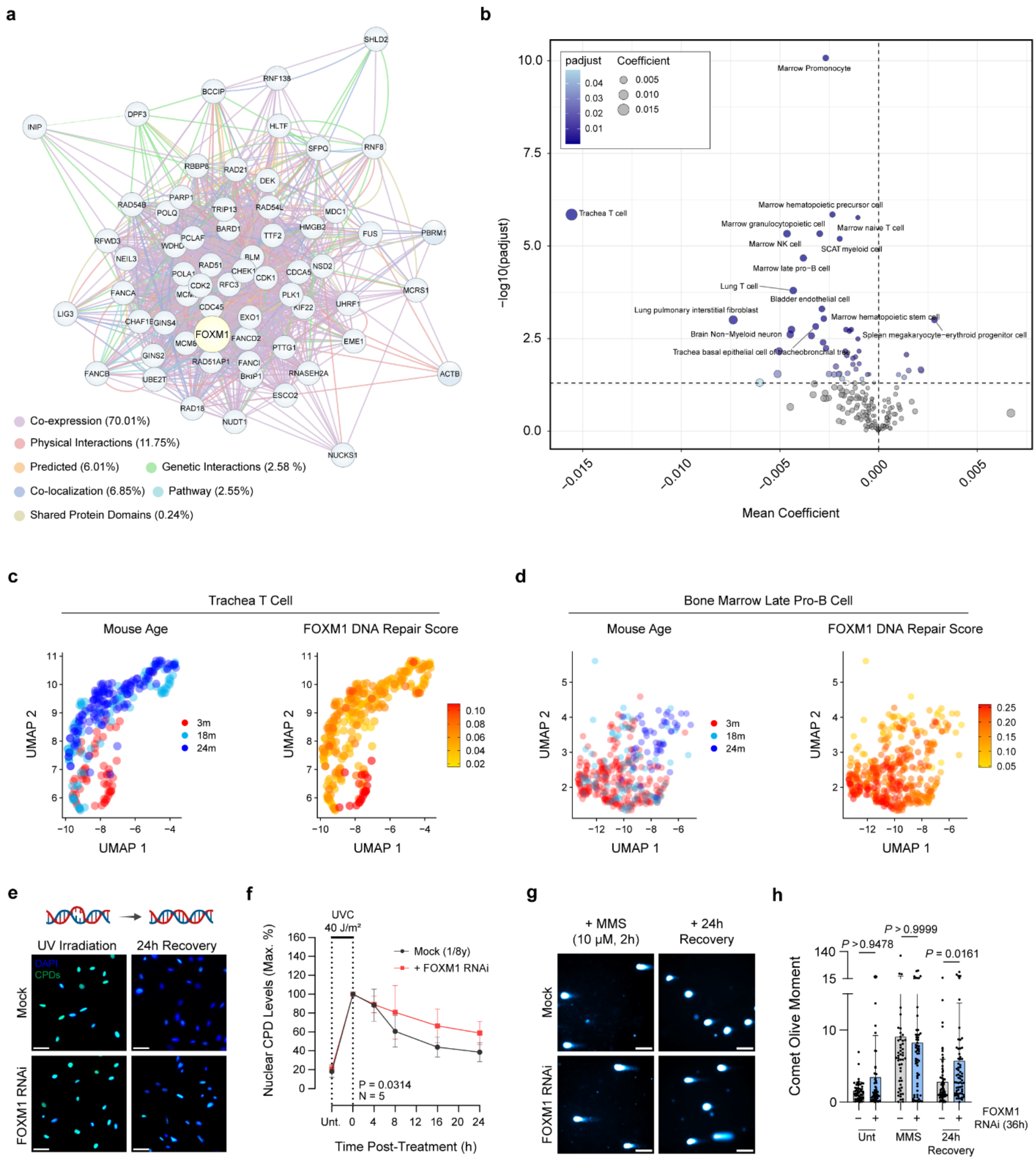


54

**SUPP. FIGURE 3**

**Supplementary Figure 3 – The FOXM1 DNA repair network is downregulated in multiple cell types during murine aging and FOXM1 repression reduces DNA repair capacity of HDFs.** **a**, Network analysis of the DNA repair genes robustly modulated by interventions targeting *FOXM1* (see Figure 2D) using GeneMANIA (Warde-Farley et al., 2010). **b**, Volcano plot of mean coefficient of variation versus -$\log_{10}$(padjust) for the FOXM1 DNA repair network in multiple cell types during murine aging. Data was obtained from *Tabula Muris Senis* (Almanzar et al., 2020). **c**, **d**, Representative UMAP plots showing age groups (in months) and signature score for the FOXM1 DNA repair network in trachea T cells (**c**) and bone marrow late pro-B cells (**d**). **e**, Representative immunofluorescence images of CPDs (green) in young HDFs (36 hr mock versus FOXM1 RNAi) immediately after UV irradiation and 24 hr following recovery. **f**, Quantification of global CPD levels in young HDFs (36 hr mock versus FOXM1 RNAi) immediately after UV irradiation and along 24 hr recovery (n = 5 independent experiments using HDFs from two distinct donors). **g**, Representative comet assay images of immediately after MMS treatment (10 µM, 2 hr) and 24 hr post-treatment. **h**, Quantification of the comet olive moment in young HDFs (36h mock versus FOXM1 RNAi) immediately after MMS treatment and 24 hr after MMS washout. Each group is represented by n ≥ 50 cells obtained from 3 independent experiments. Scale bars: 20 µm (**e**) and 50 µm (**g**). Error bars represent the standard deviation. Statistics were performed comparing indicated groups using the Kruskal-Wallis test with Dunn's multiple-comparison correction (**h**), two-way analysis of variance (ANOVA) with Sidak's multiple-comparison correction (**f**), and one-sample t-test (**b**).

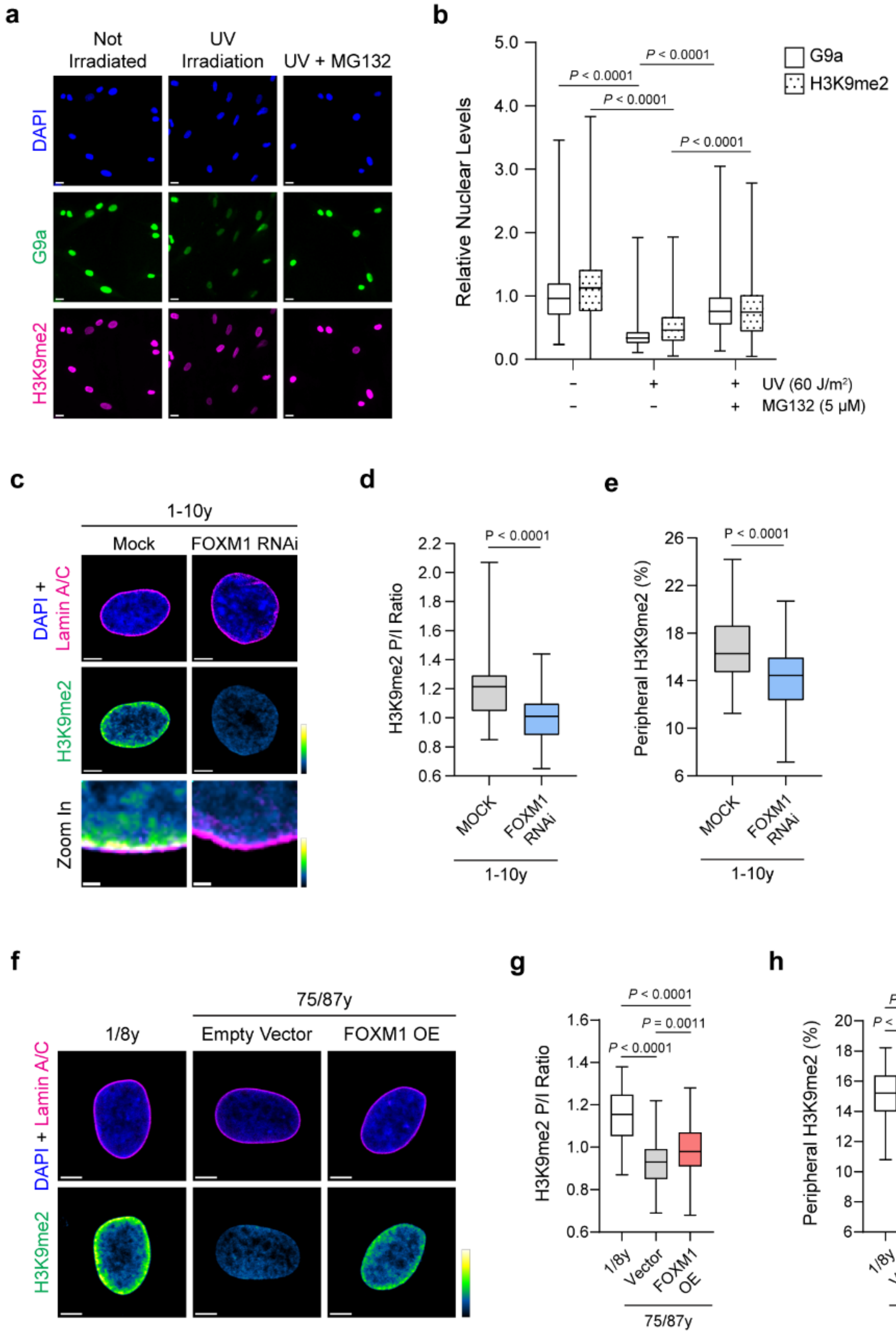


77

**SUPP. FIGURE 4**

**Supplementary Figure 4 – G9a and H3K9me2 are downregulated by multiple sources of DNA damage and can be restored in aged cells by ectopic *FOXM1* expression. a**, Representative immunofluorescence images of G9a (green) and H3K9me2 (magenta) in young HDFs either not irradiated, irradiated with UV light (60 J/$m^2$) or irradiated in the presence of MG132 (see methods for details). **b**, Quantification of G9a and H3K9me2 nuclear levels in young HDFs treated as mentioned in (a). Each group is represented by n = 900 cells obtained from 3 independent biological replicates. **c**, Representative confocal microscopy images showing H3K9me2 (green-fire-blue gradient) and Lamin A/C (magenta) immunostainings in young HDFs treated either with mock or FOXM1 RNAi. **d**, **e**, Quantification of the H3K9me2 peripheral/internal signal brightness (P/I) ratio (**d**) and percentage of H3K9me2 signal at the nuclear periphery (**e**) in young HDFs treated either with mock or FOXM1 RNAi. Each group is represented by n ≥ 68 cells obtained from 3 independent biological replicates. **f**, Representative confocal microscopy images of H3K9me2 (green-fire-blue gradient) and Lamin A/C (magenta) immunostainings in young HDFs and aged HDFs transduced with an empty lentiviral vector or overexpressing *FOXM1* (FOXM1 OE). **g**, **h**, Quantification of the H3K9me2 peripheral/internal signal brightness (P/I) ratio (g) and percentage of H3K9me2 signal at the nuclear periphery (h) in young HDFs and aged HDFs transduced with empty lentiviral vector or overexpressing *FOXM1* (FOXM1 OE). Each group is represented by n ≥ 73 cells obtained from 4 independent experiments using fibroblasts from two distinct donors. Scale bars: 20 µm (**a**), 5 µm (**c**, **f**) and 1 µm (**c**, zoom in). Statistics comparing indicated groups were performed using the Kruskal-Wallis test with Dunn's multiple-comparison correction (**b**), the Mann-Whitney test (**d**), the unpaired Student's t-test (**e**), and the one-way analysis of variance (ANOVA) with Tukey's multiple-comparison correction (**g**, **h**).

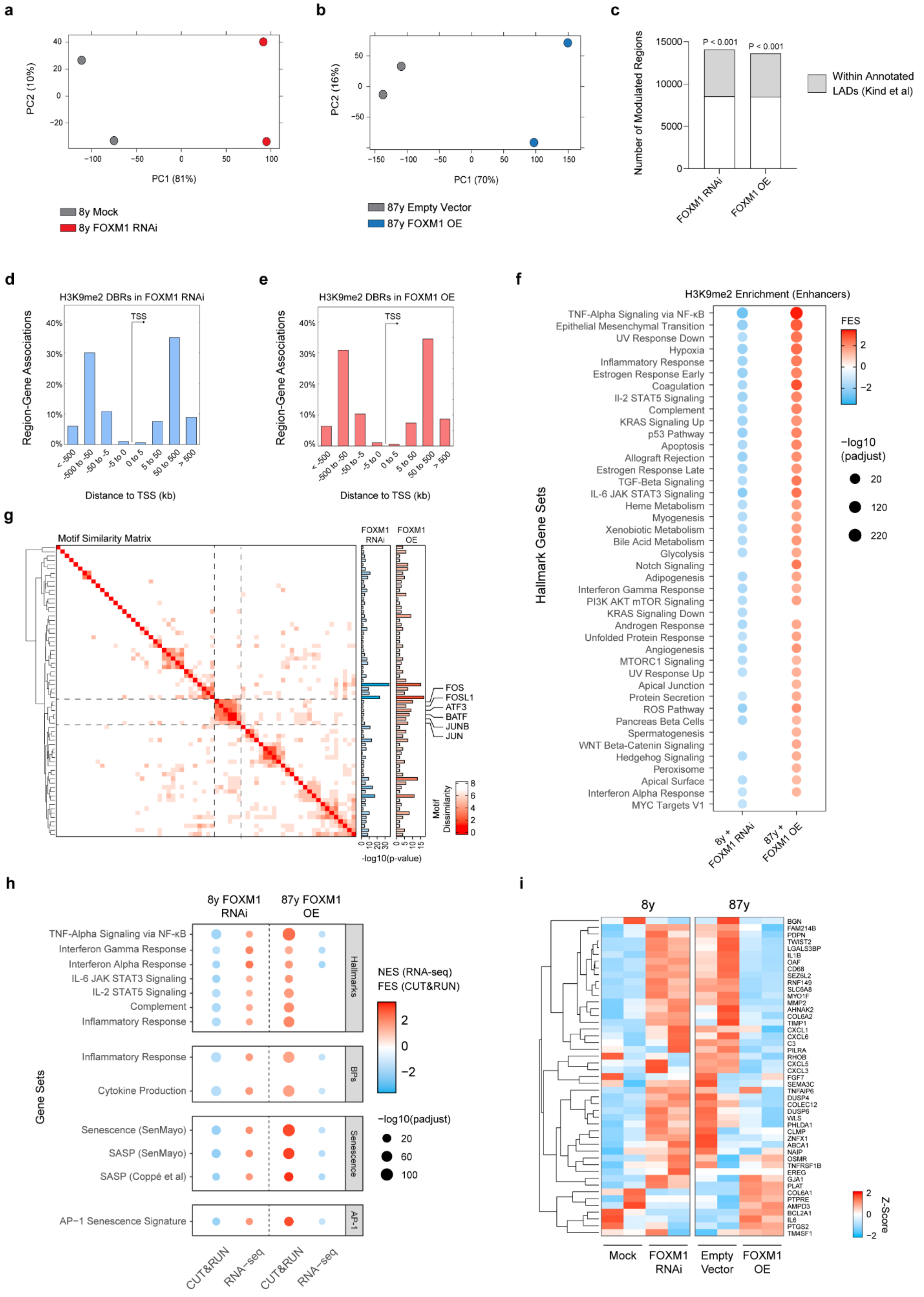




**SUPP. FIGURE 5**

**Supplementary Figure 5 – Loss of H3K9me2 is associated with upregulation of the AP-1-driven senescence signature of pro-inflammatory gene expression. a**, PCA plot of young HDFs treated with mock or FOXM1 RNAi. **b**, PCA plot of aged HDFs transduced with lentiviruses harboring empty vector or vector with a FOXM1 transgene. **c**, Overlap between regions losing H3K9me2 occupancy upon FOXM1 RNAi or gaining H3K9me2 occupancy upon FOXM1 OE and annotated conserved LADs from multiple cell types (Kind et al., 2015). **d**, Histogram of region-gene associations divided by distance to transcription start site (TSS) for regions losing H3K9me2 upon FOXM1 RNAi in young HDFs. **e**, Histogram of region-gene associations divided by distance to transcription start site (TSS) for regions gaining H3K9me2 upon FOXM1 OE in aged HDFs. **f**, Dot plot showing GREAT enrichment analysis of enhancers inside H3K9me2 DBRs for the “Hallmark” collection from MSigDB. Dots indicate gene sets enriched in regions losing H3K9me2 upon FOXM1 RNAi or gaining H3K9me2 upon FOXM1 OE in CUT&RUN data. Color gradient indicates the fold enrichment score (FES) for CUT&RUN (see Methods), and dot size indicates the $-\log_{10}$ of adjusted *p*-value. **g**, Heatmap displaying the level of similarity between the motifs (*q*-value $< 0.05$, ranked by nominal *p*-value) enriched in all H3K9me2 DBRs (i.e., without filtering for enhancer-like signatures). Bar plots on the right indicate the enrichment *p*-value after $\log_{10}$ transformation. A cluster containing motifs targeted by several subunits of the AP-1 complex is highlighted using dashed lines. Transcription factors associated with each motif in HOMER are indicated on the right of the bar plots. **h**, Dot plot showing GSEA analysis for RNA-seq and GREAT analysis for H3K9me2 DBRs (without filtering for enhancer signatures) for “Hallmark” and “Biological Processes” gene sets related to inflammation and immunity, senescence and SASP gene sets curated and validated elsewhere (Coppé et al., 2008; Saul et al., 2022), and the senescence gene signature associated with AP-1 activity (Martínez-Zamudio et al., 2020). Color gradient indicates NES (for RNA-seq data) or FES (for CUT&RUN), and dot size indicates $-\log_{10}$ of adjusted *p*-value. **i**, Heatmap showing the expression of genes in the AP-1 senescence signature (Martínez-Zamudio et al., 2020). Gene expression values are displayed after z-score transformation (higher expression in red, lower expression in blue). Statistics for sequencing data are described in detail in Methods.

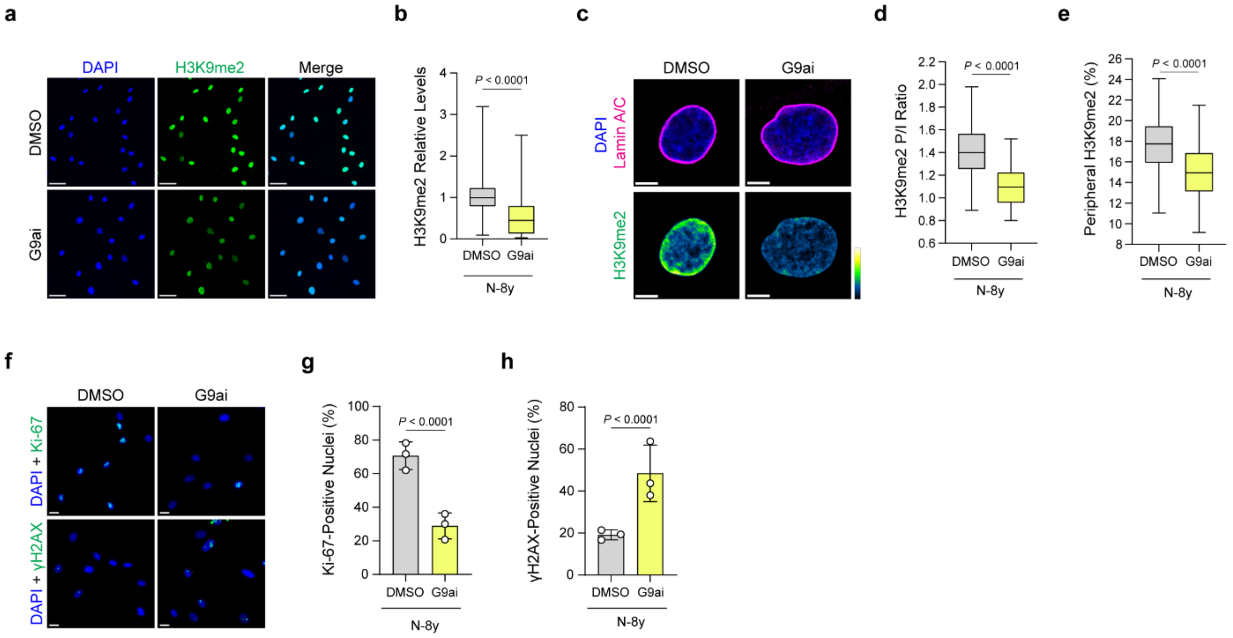


135

**SUPP. FIGURE 6**

**Supplementary Figure 6 – Inhibition of G9a activity and loss of H3K9me2 lead to senescence.** **a**, Representative immunofluorescence images of H3K9me2 (green) in HDFs derived from young donors treated with either vehicle control (DMSO) or G9a inhibitor (G9ai). **b**, Quantification of the nuclear protein levels of H3K9me2 in young HDFs treated with either DMSO or G9ai. Each group is represented by n = 900 cells from 3 independent biological replicates. **c**, Representative confocal microscopy images of H3K9me2 (green-fire-blue gradient) and Lamin A/C (magenta) immunostainings in young HDFs treated with DMSO or G9ai. **d**, **e**, Quantification of the H3K9me2 peripheral/internal signal brightness (P/I) ratio (**d**) and percentage of H3K9me2 signal at the nuclear periphery (**e**) in young HDFs treated with DMSO or G9ai. Each group is represented by n ≥ 68 cells from 3 independent biological replicates. **f**, Representative images of Ki-67 or γH2AX immunostaining (green) in young HDFs treated with either DMSO or G9ai. **g**, **h**, Percentage of nuclei staining positive for Ki-67 (**g**) and γH2AX (**h**) in young HDFs treated with DMSO or G9ai. Each group is represented by n = 1200 cells obtained from 3 independent biological replicates. Scale bars: 50 μm (**a**), 5 μm (**c**) and 20 μm (**f**). Statistics comparing indicated groups were performed using the Mann-Whitney test (**b, d**), the unpaired Student's t-test (**e**) and two-tailed Fisher's exact (**g**, **h**).
